## Supplementary Materials 1: Validations of models and task for "The successor representation subserves hierarchical abstraction for goal-directed behavior"

### S1 Appendix: Validations of task and computational models

#### TASK DESIGN

Previous investigations of community structure in statistical learning presented stimuli according to a computer-generated random walk (1–7). This ensures that the different nodes of the graph are sampled equally. Additionally, these designs often include control conditions that utilize "Hamiltonian walks", wherein graph transitions are selected by sampling each node exactly once. This procedure arguably controls for local effects of surprise associated with random walks, where between-community transitions result in higher surprise because the within-community nodes are observed more often in the recent past. When such local effects of surprise are properly controlled for, then the surprise effects associated with between-community transitions reflect "global surprise", which specifically originate from the tendency to underestimate the likelihood of such between-community transitions. Interestingly, careful analyses by (3) showed that regressing out the recency of an experienced node on response times results in virtually identical estimation of global surprise between random- and Hamiltonian walks.

Because our task design requires participants to choose freely how to walk through the graph, the sequences did not sample each node equally and no Hamiltonian control walks were enforced. In this section we provide evidence that participants nevertheless generated walks that balanced the number of visits to each node, and details on the task design that ensured this. In the section "Estimation Validation" we provide evidence that the design yields separable local- and global surprise effects.

##### Training phase design

In the training phase of the task, each participant searched for a goal painting 75 times, i.e. they performed 75 "miniblocks". Each of the 15 rooms was presented as the goal room (and consecutively as the start room) exactly 5 times. Main text Fig. 1A shows how different rooms can lie at the "boundary" of two wings, allowing for transitions between two wings (i.e. rooms 1, 5, 6, 10, 11, 15). The remaining rooms only connect to rooms in the same wing. We will refer to these respectively as "boundary rooms" and "deep rooms". These differences imply different distances between starting locations and goal locations, each with their own expected value. Fig. S1 illustrates all different possible start-goal combinations. If start and goal locations would have been randomly sampled, the expectations of sampling these different distances are listed in Fig. S2. For every participant prior to the training phase, we sampled sequences of 75 goals until we found a sequence that matches these expectations for room identity and goal distance exactly.

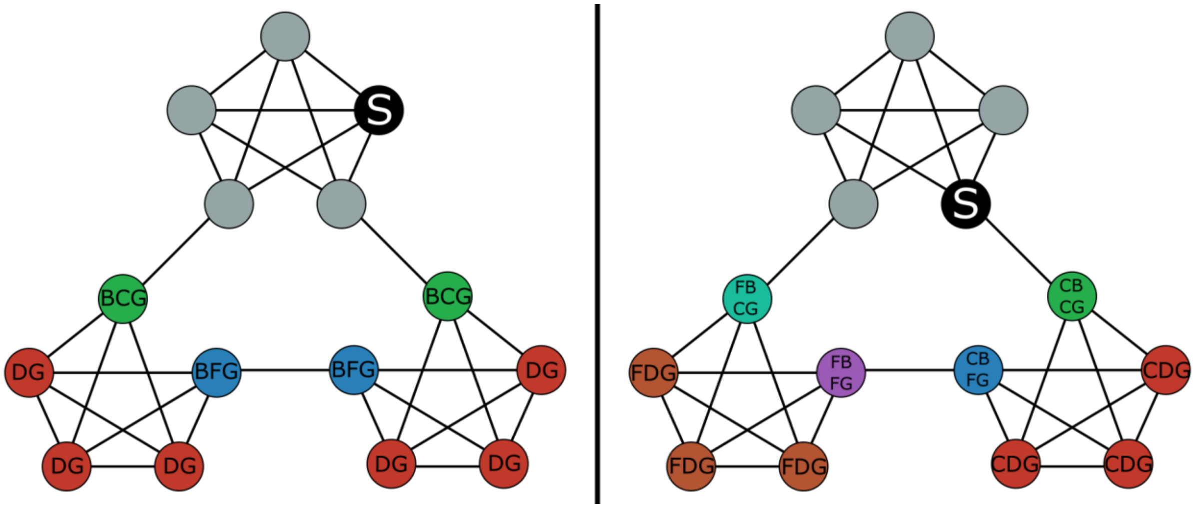


**Fig S1 Illustration of possible start-goal combinations.** The starting room is indicated as a black filled node with an "S". Only rooms outside of the starting wing are considered as goals. An asymmetry in the two possible goal wings is introduced if the starting room is a "deep" room, meaning a room not connected to any room in another wing (left panel), vs. a "boundary room", meaning it has a connection to a room in an adjacent wing (right panel). If the start location is a deep room, then the goal location can be a deep room in a different wing, or it can be a boundary room in a different wing. In the left panel, deep goals are marked with "DG". If the goal is a boundary room in a different wing, then it can be a boundary room that is directly connected to the starting wing ("boundary-close goal" or "BCG"), or it can be a boundary room that is not connected to the starting wing ("boundary-far goal" or "BFG"). Note that for deep start rooms, the start-goal distances are symmetric across the two possible goal wings. By contrast, if the starting location is a boundary room, then the goal location can be in the wing that is directly connected to that starting room, or in the wing that is not connected to that starting room. This is indicated by prefixing the goal respectively with "close" or "far". In the right panel, "close deep goals" are marked as "CDG" and "far deep goals" are marked as "FDG". Further, "close boundary-close goals" are marked as "CBCG", "close boundary-far goals" are marked as "CBFG", "far boundary-close goals" are marked as "FBCG", and "far boundary-far goals" are marked as "FBFG".

| start → goal | p(start) | p(goal\|start) | p(start, goal) | $\mathbb{E}$ over 75 trials |
|---|---|---|---|---|
| Deep → Deep | 3/5 | 3/5 | 9/25 | 27 |
| Deep → Boundary-Close | 3/5 | 1/5 | 3/25 | 9 |
| Deep → Boundary-Far | 3/5 | 1/5 | 3/25 | 9 |
| Boundary → Close Deep | 2/5 | 3/10 | 3/25 | 9 |
| Boundary → Far Deep | 2/5 | 3/10 | 3/25 | 9 |
| Boundary → Close Boundary-Close | 2/5 | 1/10 | 1/25 | 3 |
| Boundary → Close Boundary-Far | 2/5 | 1/10 | 1/25 | 3 |
| Boundary → Far Boundary-Close | 2/5 | 1/10 | 1/25 | 3 |
| Boundary → Far Boundary-Far | 2/5 | 1/10 | 1/25 | 3 |

**Fig S2 Start-goal expectations.** Table indicating the expected selection of each start-goal identity, both in exact proportion ("p(start, goal)") and in exact number when considering a total of 75 miniblocks ("E over 75 trials"). The proportions can be understood by looking at the graph in Fig. S1. For p(start), it is the fact that there are always 2 boundary rooms and 3 deep rooms (for a total of 5 rooms in total) in each wing. p(goal|start) can then be obtained by assuming a particular room was selected as the start, and considering the 10 possible goal-rooms in the other two wings. Multiplying "p(start)" with "p(goal|start)" gives the joint probability of a particular start-goal combination ("p(start, goal)"). Multiplying this by 75 gives the expected number of times this start-goal combination is encountered over 75 miniblocks ("E over 75 trials").

#### Action-outcome mapping and node visit expectations

In this section we provide the rationale behind the "preferred" and "non-preferred" transitions as described in the main text, by showing how alternative task designs fail to yield a balanced sampling of all nodes. It is important to note that there are two components that determine the sequence of nodes experienced by a participant. The first is the action-outcome mapping, and the second is the behavioral policy applied by the participant. We do not control the behavior of the participant. Hence, we cannot manipulate this component in our experimental design to guarantee a balanced sampling of all nodes. We assume that the behavior of the participants can be approximated as either a "stochastic" policy, meaning the participants choose a random action on each new step in the task, or a "one-direction" policy, meaning the participants select only one action over the course of the entire task. If the sampling of nodes is shown to be uniform over the course of the experiment under these policies, it would also be uniform under random mixtures of these policies. Crucially, participants that switch between bouts of repeatedly selecting <z> and repeatedly selecting <m> dependent on the current goal (which is randomly determined), occasionally experiencing attentional lapses yielding unstructured stochastic behavior, should yield behavioral policies that correspond to the distribution of node visits as simulated under these policies.

We wish to validate that sequences of experienced states generated by these reasonable default policies do not differ from sequences generated by true random walks as leveraged in previous experiments (1–7). Behavioral policies are defined with respect to the selection of the different actions (i.e. pressing the <z> and <m> key) and are not under experimental control. By contrast, the action-outcome contingencies for each state are under experimental control. We can quantify how uniform the sampling of nodes is under a particular action-outcome mapping given a particular behavioral policy, by computing the proportional difference between the number of visits to the most and the least visited nodes (the "visit asymmetry"). Additionally, we can compute the number of transitions before leaving a particular community (the "dwell time"). As a benchmark, we compute these summary statistics for sequences generated from a true random walk in keeping with previous experiments (1–7). We aim to find an action-outcome mapping that matches these summary statistics for the "stochastic" and "one-direction" policies. Note the key difference between the benchmark random walk and the stochastic policy is that the latter randomly selects an action, after which a transition is generated according to the relevant action-outcome mapping, whereas the former directly generates transitions from a uniform distribution over relevant edges without concern for any action-outcome mapping. Below, we consider several reasonable action-outcome mappings and show how they fail to match a random walk, except for our final "balanced" action-outcome mapping (main text Fig. 1B). For each policy and action-outcome mapping, we generate sequences according to the rules of the training phase, i.e. 75 miniblocks that balance the selection of goals and start-goal combinations. Simulations presented here thus simultaneously validate our action-outcome mapping, and the design of our training phase, when compared with sequences generated under a benchmark random walk.

#### Arbitrary action-outcome mapping

The first action-outcome mapping one could design, is to randomly assign two of four possible outcomes of each node to action (i), and the other two outcomes to action (ii). This can be done in advance, separately for each node. This way, there is a high demand on the participants to learn specific action-outcome contingencies. Note that no meaningful hierarchical choice policy exists under this action-outcome design, however response times can still reflect community structure. In a pilot study, participants exposed to these action-outcome contingencies reported high levels of frustration and generated unnecessarily long sequences to reach goals. Because consistently pressing the same key does not always result in between-wing transitions for this set of action-outcome mappings, simulated (and real) agents that apply this policy can get "stuck". Therefore, one-direction policies are not simulated for this action-outcome mapping. A stochastic policy for this arbitrary action-outcome design will yield sequences with identical properties to a stochastic policy under the "consistent" action-outcome mapping (see below) and in fact these are both mathematically identical to the benchmark random walk we simulated as a benchmark. We only report simulations of the stochastic policy for the "consistent" action-outcome mapping as a proof of concept and omit the arbitrary action-outcome mapping to preserve space.

##### Consistent action-outcome mapping

In order to elicit a hierarchical choice policy that would be meaningful to the participants, we expected that consistently taking the same action could intuitively correspond to traveling in one direction through the museum. Hence, one action should "rotate" in one direction through the different wings of the museum, whereas the other action should rotate in the opposite direction, resulting in the design illustrated in Fig. S3. Here, green and pink arrows correspond to transitions that move or fail to move in the intended rotational direction. Blue arrows are equally consistent with the intended rotational direction. Note that not every node receives the same amount of incoming arrows, which yields visit asymmetries under a "one-direction" policy compared to a benchmark random walk (Fig. S4, row 3 columns 1 and 2). Additionally, note that the nodes from which the blue arrows project do not allow for within-wing transition back to this state and are thus excluded from the one-directional sequences starting from other nodes in this wing. Further, observe that the "boundary" node allowing for the intended between-wing transition has an outgoing pink arrow with a reciprocal green arrow pointing directly back at it. This allows for a "direct recovery" from an unintended transition away from the desired between-wing transition. This yields a substantially shorter within-wing dwell time under a one-directional policy compared to a benchmark random walk (Fig. S4, row 3 column 3). Additionally, it contributes further to the visit-asymmetry, preferring the node on which the green arrows "converge", and yielding successively less visits to nodes "further away" from it when following the pink arrows.

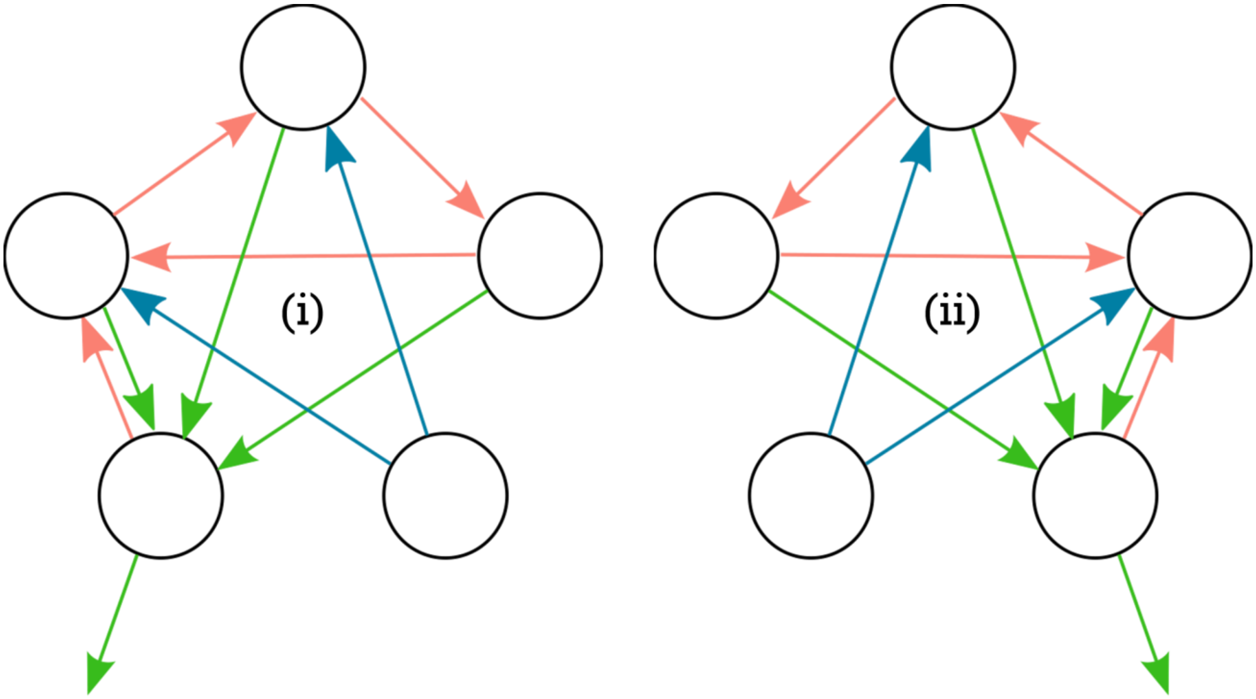


**Fig S3 Consistent action-outcome mapping.** Prototype for a "naïve" mapping that allows one action to correspond to a "rotation" through the wings of the museum in a particular direction. Green arrows are outcomes that correspond to this intended rotation, while pink arrows indicate "unintended" outcomes of these actions. Blue arrows both equally correspond to transitions in agreement with this intended rotation.

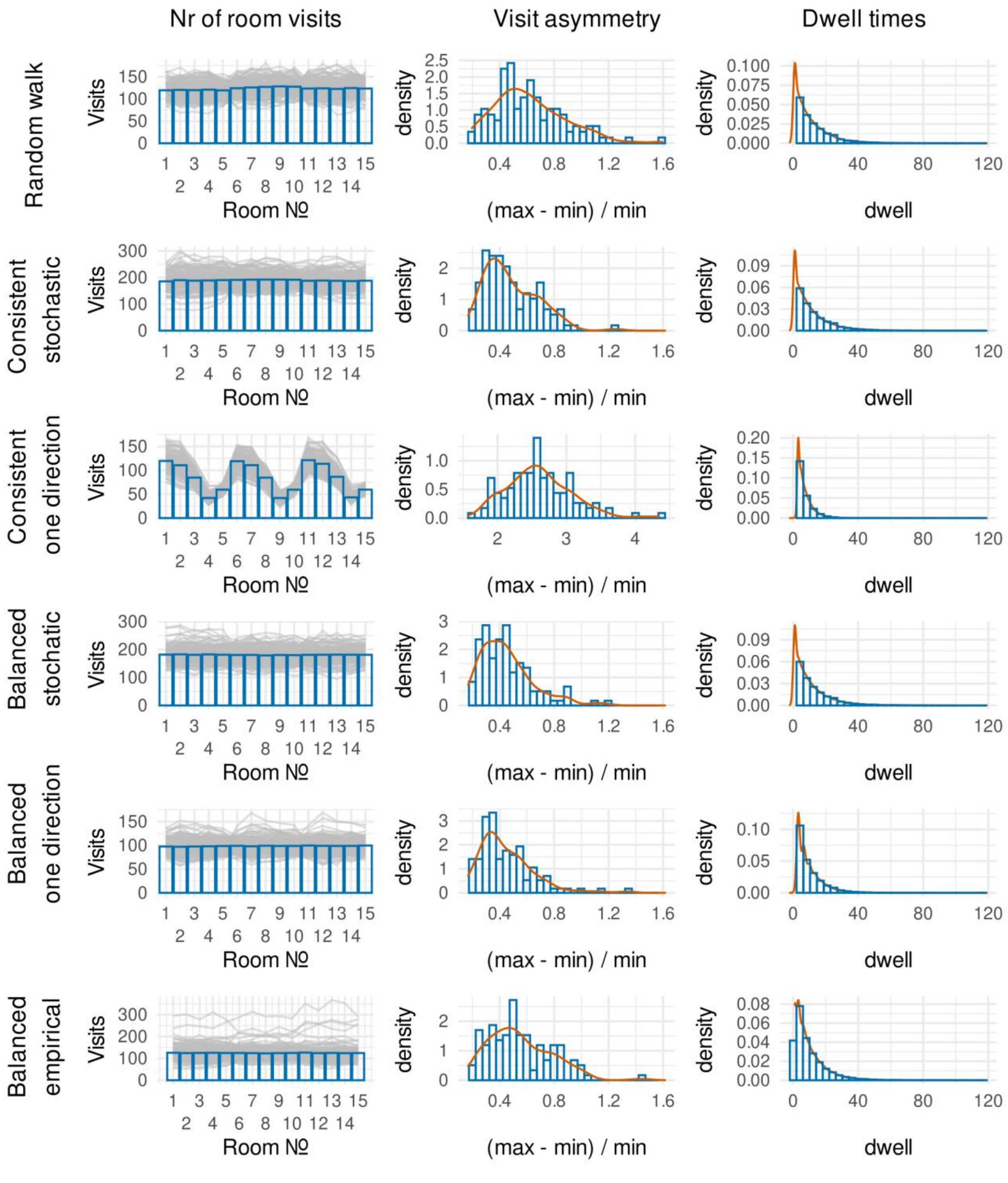


**Fig S4 Node visit expectations for different action-outcome mappings and behavioral policies.** We compare the transition statistics (columns) of simulations of different combinations of action-outcome mappings and behavioral policies (rows). The first column plots the expected number of visits to each node. The second column plots the "visit asymmetry", quantified as the proportion of visits by which the most visited node outweighs the least visited node. The third column plots the "dwell time", the expected number of transitions made within a community before leaving it. For the combinations of action-outcome mappings and behavioral policies, each row is labeled by first indicating the action-outcome mapping ("balanced", main text Fig 1B, or "consistent", Fig. S3) and then the behavioral policy ("stochastic" or "one direction", see body text). Additionally, we consider a benchmark "random walk", and the statistics for our empirically collected data ("balanced empirical"). For the simulation of behavioral policies, we generated 120 artificial agents completing the training phase of the museum task. For the benchmark random walk we drew 1850 transitions from an undirected graph as shown in main text Fig. 1A. Note that the node visits (first column) appear to be biased for the simulated combination of consistent mapping and one direction policy ("consistent one direction", row 3). This simulation also yields a substantially higher visit asymmetry (second column); note that the x-axis is plotted on a different scale compared to the other simulation. Finally, the "consistent one direction" combination yields a dwell time about half that of all other simulations.

#### Balanced action-outcome mapping

We identified the dwell times and visit asymmetries generated by the consistent action-outcome mappings as problematic, given that under a one direction policy the expected behavior strongly deviated from that of the benchmark random walk (Fig. S4). If specific nodes were expected to be visited more often than others, this would confound our ability to make inferences about the community structure. Additionally, shorter dwell times within a community would make the community structure effect less salient to the participants. We therefore considered an alternative mapping that maintained the rotational "hierarchical" mapping across wings, but yielded sequences that displayed similar visit asymmetry and dwell time distributions as the benchmark random walk. Taking the "consistent" action-outcome mapping as a starting point, we identified two key problems. Firstly, the node out of which the blue arrows project is not included as a within-wing outcome (Fig. S3). However, this node should participate in the within-community sequences just as much as all other nodes. Secondly, the "direct recovery" from the unintended pink arrow out of the boundary node is responsible for yielding shorter dwell times and asymmetric node visit expectations. We addressed these two concerns jointly by removing the "direct recovery" green arrow, and replacing it with an arrow pointing toward the other boundary node (out of which the blue arrows project). Additionally, we reassigned the blue arrow projecting back to this node so that each node has exactly two incoming arrows for each action. The final "balanced" mapping (main text Fig. 1B) yields dwell times and visit asymmetries that are very comparable to a benchmark random walk, validating both this action-outcome mapping and the design of the training-phase as yielding unbiased walks through the graph, for both the stochastic (Fig. S4, row 4) and one direction (Fig. S4, row 5) policies. Crucially, the empirical data collected under this action-outcome mapping (Fig. S4, row 6) are highly comparable to those of the benchmark random walk.

As noted in the main text and main text Fig. 1C, this new action-outcome mapping results in some states leading to the same outcome for both actions. For example, state 1 leads to state 2 (main text Fig. 1A) with 50% probability irrespective of which key the subject presses. We refer to these as "preferred" transitions. These preferred transitions result in some nodes being more often followed by a specific other node. Crucially however, it is not the case that some nodes are visited more often than other nodes overall, as evidenced by the visit asymmetry shown in Fig. S4 (rows 4 and 5, column 1). However, this does yield differences in the successor representation prediction error for preferred transitions compared to non-preferred transitions. Specifically, because successor representations are policy-dependent, successor vectors should be biased to represent preferred outcomes, thus yielding smaller prediction errors upon preferred transitions. Beneficially, this increases the (within-community) variance that could be captured by a successor representation model, both with respect to state prediction errors and expected values. Meanwhile, the between-community effect can still be computed in an unbiased manner (see section "Modularity Measurement"). Any other systematic influences can be controlled for by considering preferred outcomes as a covariate in our regression models (see main text methods).

### COMPUTATIONAL MODELS

We simulated the museum task using each of the four cognitive models described in the main text: null, model-based ("MB"), explicit hierarchical ("Exp"), and successor representation models. To approximate behavior under a null model, we simulated both stochastic ("Rand") and one-direction ("One") policies to indicate baseline performance. For the successor representation model we consider two agents, one with the discount factor value set to 0.1 ("SR01") and another with the discount factor value set to 0.9 ("SR09"). This leads to 6 simulated agents in total. For each agent, we simulated the performance in 120 instances of the task, with the "preferred" action-outcome contingencies, for both the training and the test phases. Fig. S5 (top) shows the total reward accumulated for each agent. For the model-based agent, we ran a value-iteration algorithm to convergence for every possible goal before executing the task, and had the agent select the action with the highest value in each case. For the explicit hierarchical model, the agent selected the correct "rotational" action whenever it was in a different wing from the goal wing. Within the goal wing, the agent selected the optimal model-based action. The successor representation was learned using temporal difference learning and used to compute action-values (see main text methods). It always selected the action with the highest estimated value, or a random action if values were identical.

Fig. S5 (top) shows the model-based agent performs the best (as defined by total reward), as expected because it follows the optimal policy for every possible goal. The figure also illustrates that a higher discount factor for the successor representation leads to slightly better performance relative to the lower discount factor, potentially linked to the faster updating of the task model. As expected, the explicit hierarchical agent yields good performance, but accumulates a lower total reward than the model-based agent. The null agents perform much worse than the other agents.

Fig. S5 (bottom) shows matrices visualizing the action policies for the model-based, explicit hierarchical, and successor representation agents (both "SR01" and "SR09"). Null agents are omitted since their policies are not dependent on states or goals. The matrices are organized by states on the y-axis and goals on the x-axis (both numbered 1-15 as in the main text Fig. 1A). Colors indicate the proportion with which a particular action was selected for that state-goal combination over the course of all simulations of that agent. More blue squares indicate higher proportion of action (i), and more orange squares indicate a higher proportion of action (ii) (actions labelled as in main text Fig. 1B). Yellow squares indicate a uniform distribution of selected actions. Especially when looking at the explicit hierarchical policy, the matrix can be seen to have a block structure. This policy always selects the same action for all states in the same community, for all goals in a specific other community. We highlight these communities with pink and green squares around the blocks, green indicating that (i) is the optimal rotational action, and pink indicating that (ii) is the optimal rotational action. Interestingly, the model-based (optimal) policy seems to select different actions for different states in the same community, and this also depends on different goals in the same goal community. Only the successor representation models yield squares that interpolate between orange, yellow and blue, since it does not follow a precomputed deterministic policy. In some states, which action is considered more optimal is dependent on the recent transition history, since the successor representation is constantly being updated using the temporal difference learning rule. Interestingly, more yellow squares appear in the low discount factor ("SR01") case, than in the high discount factor ("SR09") case. Instead, the higher discount factor more closely approximates the true model-based agent, which follows the optimal policy. This indicates the higher discount factor agent is faster to adapt to the currently relevant goal, hence yielding higher total reward. Overall, these simulations indicate the different models correspond to different choice behavior, and that the discount factor parameter of the successor representation “tunes” the accuracy of the behavior of the agent.

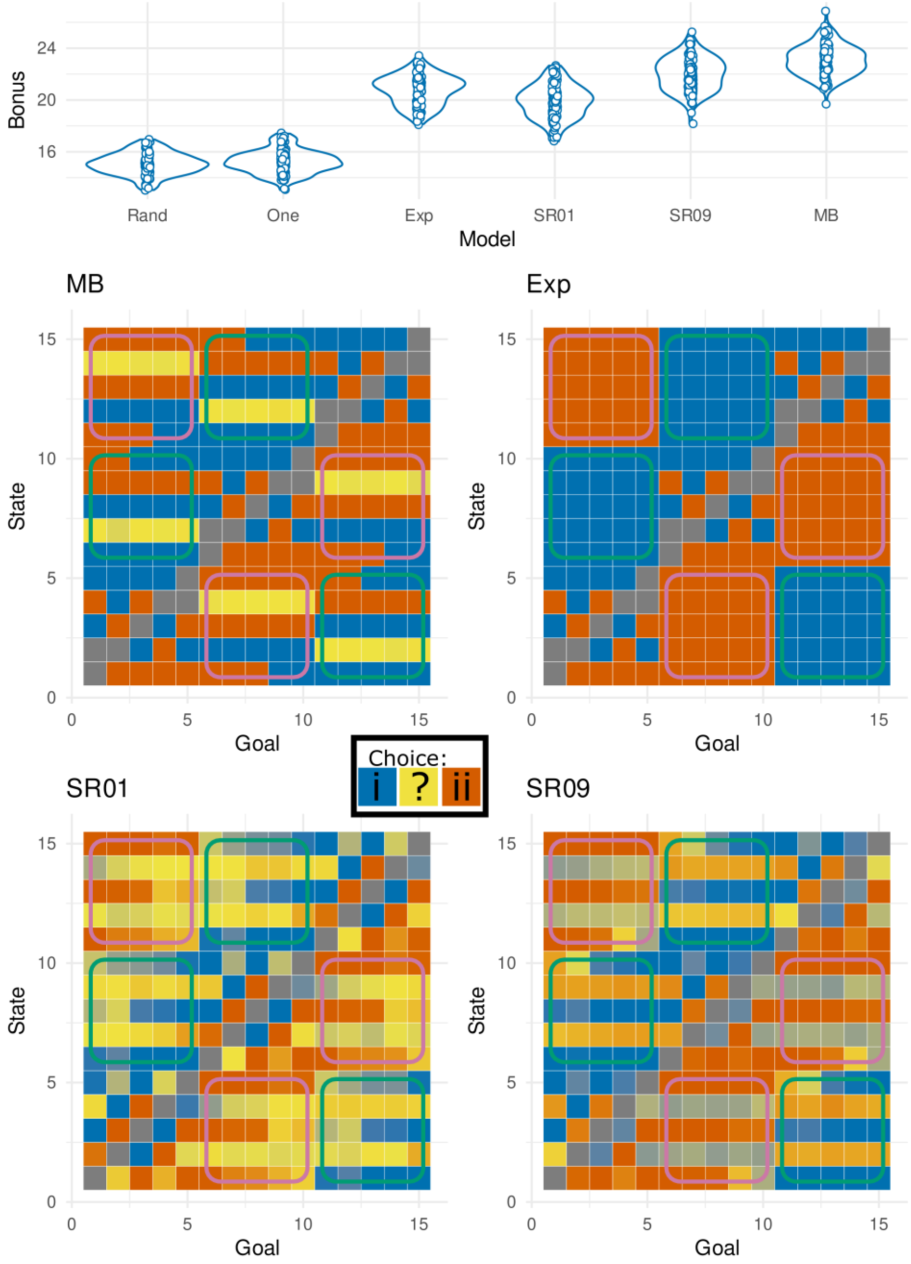


**Fig S5 Simulations of accumulated rewards and policies of different agents.** Top figure shows the total accumulated rewards for 120 agents of each model in the museum task. Bottom 4 matrices show greedy choice policies for the model based (MB), explicit hierarchical (Exp), and the successor representation (SR) models. The successor representation was simulated twice, once with a discount factor of 0.1 (SR01) and once with a discount factor of 0.9 (SR09). Matrices reflect all different goals (x-axis) and all different state occupations (y-axis) with states labeled as in main text Fig. 1A. Blue squares correspond to action (i) and orange squares to action (ii). Yellow squares indicate identical values or choice-proportions near 0.5. Superimposed green and violet borders are used to indicate rotational symmetry between start and goal wings. Green borders correspond to rotational symmetry where action (i) leads to the goal wing. Violet borders correspond to wings where action (ii) leads to the goal wing.

#### Parameter and model recovery

In order to validate the cognitive models estimated from our data, we simulated 200 individual participants from each cognitive model. For each simulated participant we randomly (with replacement) drew a participant from our real data set and used their sequence of states and actions to provide realistic task experience from which to generate artificial data. All parameter values were randomly generated from the prior distributions as specified in the main text methods. We fitted all 4 models to each simulated participant, however instead of full MCMC we relied on variational approximation as implemented in Stan. While variational approximation is biased and not recommended for inferences based on parameter estimation (8), it has been shown as an efficient technique for the validation and comparison of computationally expensive cognitive models (9). It yields a posterior predictive distribution that can be used to compute the approximate leave-one-out cross-validation score and the pseudo-BMA+ weights similar to full MCMC. Since variational inference is called on the same model code as our main MCMC estimations, good performance with variational inference provides a lower bound for the performance of our main MCMC estimates.

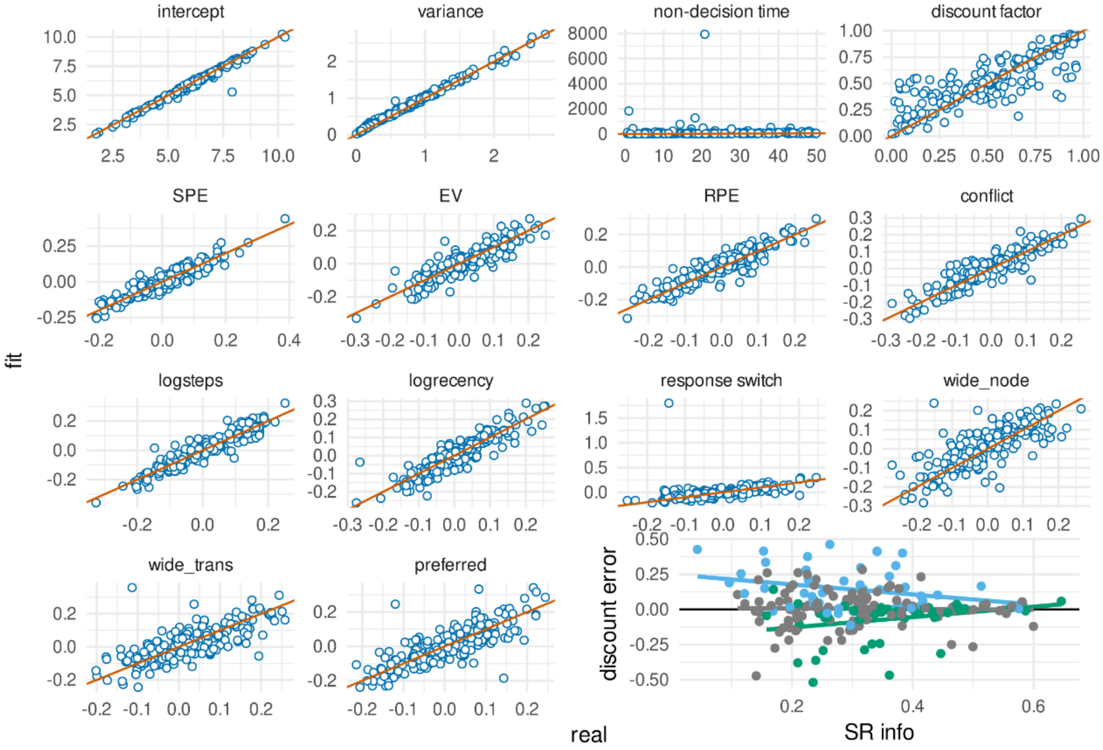


**Fig S6 Parameter recovery for the response time successor representation model.** Panels show the ground-truth generated value ("real", x-axis) plotted against the posterior mean of the model fitted to data generated using these ground-truth parameter values ("fit", y-axis). The orange line indicates matching values between the real and fitted values. Parameters include the intercept (top left), the error variance (top: second), the non-decision time (top: third) and the discount factor (top right). Regressor coefficients include the successor representation derived parameters (SPE: state prediction error, EV: expected value, RPE: reward prediction error, conflict; second row). Additionally, nuisance regressors related to task execution (logsteps: number of steps on current miniblock, logrecency: number of trials since last encounter of current state, response switch: binary indicator of changed response key) and nuisance regressors related to the action-outcome contingencies (see main text Fig. 9; wide node, wide trans, preferred). The panel on the bottom right shows the error of the recovery of the discount factor, plotted against the sum of the absolute values of the four successor representation regressor coefficients (SPE, EV, RPE, and conflict). Blue dots represent ground-truth discount factors below

0.25, while green dots represent those above 0.75. It can be seen these errors shrink to 0 as the discount factor becomes more informed, effectively “washing away” the regularization from the uniform prior.

Parameter recovery for the response time successor representation model is shown in Fig. S6. Parameter recovery for the other models is appended at the end of this document (Fig. S14). For the successor representation, it can be seen the discount factor estimation is somewhat noisy (top right panel). Specifically, “real” values (x-axis) further away from 0.5, appear regularized toward a fitted value (y-axis) of 0.5. This is because we report only the posterior means of the fitted models and our Beta(1,1) prior has a posterior mean of 0.5. The discount factor can only be estimated accurately if the regressors associated with the successor matrix have a strong influence on the response time. Adding together the absolute values of these regressors (SPE, EV, RPE, conflict) gives us a measure of “successor representation information” (SR info). In the bottom right, we plot the error in the estimation of the discount factor against this SR info. For the purpose of illustration, dots are colored blue for real discount factors below 0.25, and green for real discount factors above 0.75. It can be seen values further away from 0.5 are increasingly estimated more accurately when more SR info is available, visualized by regression lines of the corresponding color.

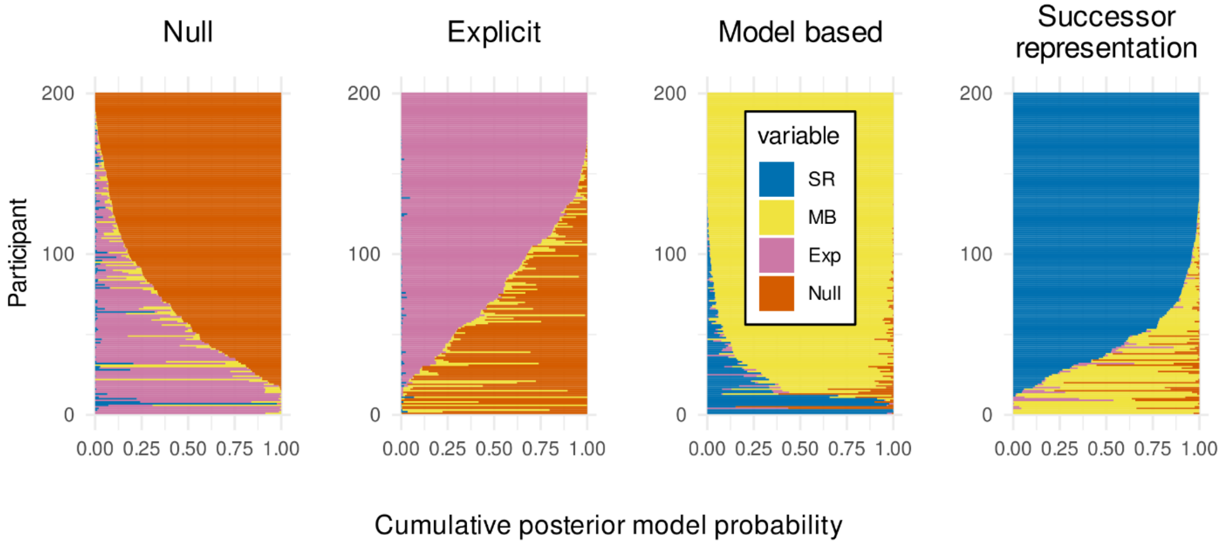


**Fig S7 Model recovery for the response time models.** Figures show horizontally stacked bar charts visualizing the posterior model probabilities for our 200 simulated participants under each ground-truth model. Posterior model probabilities derived from random effects Bayesian model comparison between four cognitive models, “Null” (orange), explicit hierarchical “Exp” (pink), model-based “MB” (yellow), and successor representation “SR” (blue). The plots are titled by ground-truth model used to generate data. Participants were sorted by posterior model probability for the ground-truth model.

Fig. S7 shows the posterior model probabilities for all 200 simulated participants for each ground-truth model. Additionally, we randomly subsampled 120 out of 200 simulated participants to match our real dataset size and computed a Bayesian omnibus risk (BOR) value. We did this 100 times, and observed all BOR < $10^{-8}$ in favor of the ground-truth (all pxp > 0.955). Interestingly, it can be seen the two “simpler” models (the null and the explicit hierarchical model) are more often confused with each other (pink vs. orange), while the more “complex” models (the model based and the successor representation model, which both contain a discount factor parameter) are also more confused with each other (yellow vs. blue). Overall, recovery of these more complex models appears more consistent. This is generally the case, because model comparison procedures tend to favor more complex models over simpler ones. It is thus important to note that the recovery of the null model was significant in all our simulations, indicating our main results are not due to a preference for more complex models. The fact that at the individual level, the data generated from the null model virtually never shows any evidence in favor of the successor representation, provides strong evidence that the complexity of the successor representation model is properly penalized by our model comparison technique.

We apply the same simulation and recovery procedure to the models of choice data and find good recovery of both the models and the parameters (Fig. S8). Again every group-level model comparison is significant (all BOR < $10^{-11}$) in favor of the ground-truth model (all pxp = 1).

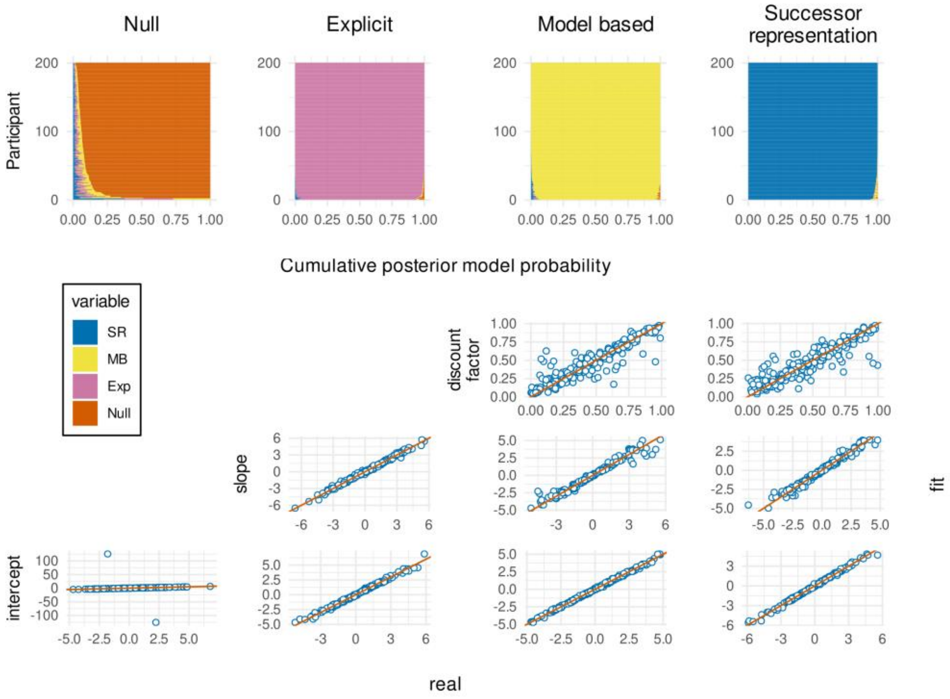


**Fig S8 Model and parameter recovery for the choice models.** Top row: Figures show horizontally stacked bar charts visualizing the posterior model probabilities for our 200 simulated participants under each ground-truth model. posterior model probabilities derived from random effects Bayesian model comparison between four cognitive models, “Null” (orange), explicit hierarchical “Exp” (pink), model-based “MB” (yellow), and successor representation “SR” (blue). The plots are titled by ground-truth model used to generate data. Participants were sorted by posterior model probability for the ground-truth model. Bottom: parameter recovery for each ground-truth model, placed in the same column as the posterior model probabilities. Panels show the ground-truth generated value (“real”, x-axis) plotted against the posterior mean of the model fitted to data generated using these ground-truth parameter values (“fit”, y-axis). The orange line indicates matching values between the real and fitted values. The null model only contains an intercept. The explicit model additionally contains a regression coefficient for the correct “rotation” (see main text methods). The model based and successor representation models contain a regression coefficient for the expected values of the different actions, which additionally depend on a discount factor parameter (see main text methods).

#### ESTIMATION VALIDATION

We aim to validate that our model fits yielded successor representations and community structure effects that were both stable and interpretable. We first provide details on the sources of state prediction errors to show that the between-community transitions yield response slowing irrespective of local (i.e. non-SR related) surprise signals, also termed "recency" effects. We then provide evidence that the regressors derived from the successor representations are relatively stable over the course of the testing phase, suggesting that the successor representation matrices were learned appropriately during the training phase.

#### Global and local surprise

Our model of the successor representation included a "nuisance" regressor capturing the log-transformed recency, i.e. the number of trials that have passed since the current state was last encountered. It has previously been shown that including such a recency regressor yields an estimated between-community slowing effect about the same value as what is obtained for a Hamiltonian walk, which experimentally controls for recency (3). In other words, global surprise as derived from the communities is a separable effect from the local surprise dependent on the recent history of state visits. The community slowing effect in our task is captured by the state prediction error derived from the successor representation. Because we estimated our models using MCMC, it is possible to inspect the correlation between the samples of the coefficients for the recency regressor and the state prediction error. Fig. S9 plots for all samples the regression coefficient of the log-recency against the regression coefficient of the state prediction error and shows lines fitted to each of the 120 individual participants. If local and global surprise were problematically confounded in our analysis, we would expect a majority of downward lines to be salient, indicating a trade-off between the two coefficients. Instead, we observe mostly flat lines, indicating no relevant overlapping variance in the estimation of recency and between-community slowing effects.

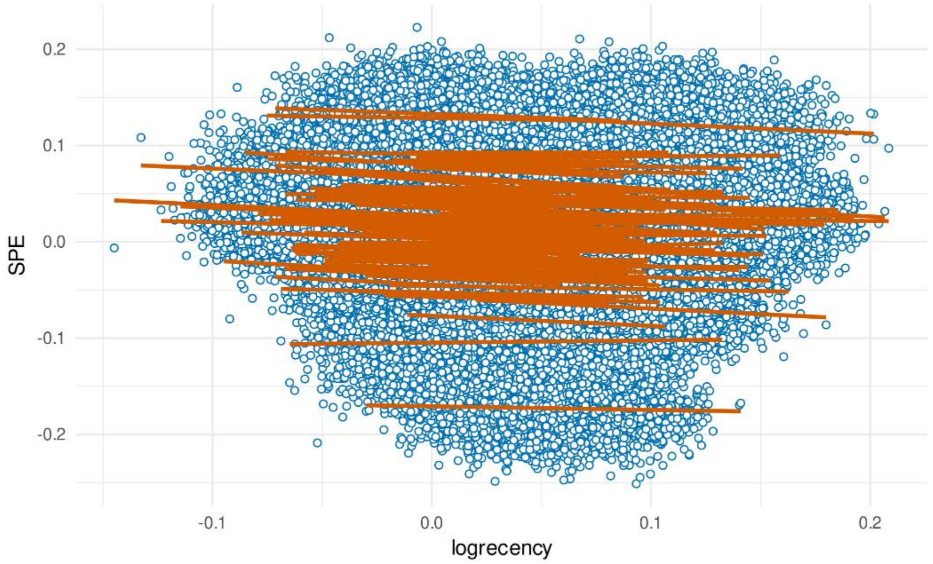


**Fig S9 Correlation between log-transformed recency and state prediction error coefficients.** Blue dots represent single MCMC samples across all participants. Orange lines are fitted to the samples of each separate participant. Problematic correlations between local and global surprise signals dictate a trade-off between the recency and the state prediction error regressors leading to negative slopes of the orange lines. No such trade-off is observed.

#### Learning convergence of the successor representation

We confirmed that the fits of the successor representation model converged based on the Gelman-Rubin statistic and the absence of divergences. Another important question regarding convergence concerns the stability of the successor representation during the testing phase. The successor matrix starts out as an identity matrix and incorporates more structure when accumulating experience. This means state prediction error will be initially high, and expected value initially low. After sufficient experience, the successor matrix will reflect an appropriate model of the task, and no further structure can be incorporated except for local variations in policy based on the changing goals. This means the values of the regressors derived from the successor representation will fluctuate around an average value, instead of show an increasing or decreasing trend over the course of successive trials. Fig. S10 plots the regressor values as derived from the maximum a-posterior fit for each participant over the course of the testing phase. Orange lines represent generalized additive models fit to the time course of individual participants. Lack of overall trends confirms the absence of any such trends in the testing phase, indicating the initial learning successfully converged in the training phase. By contrast, Fig. S11 plots how these regressor values would be assigned during the training phase. It can be seen the state prediction errors start out high early in the training phase, and reach asymptote after around 1000 trials, indicating learning has converged. For the value driven effects, it can be seen the mean is relatively stable throughout the training phase, but the variance increases with more experience. As the successor representation model gains more experience, it is able to make stronger predictions about upcoming states, and thus assign higher (and lower) values and associated prediction errors.

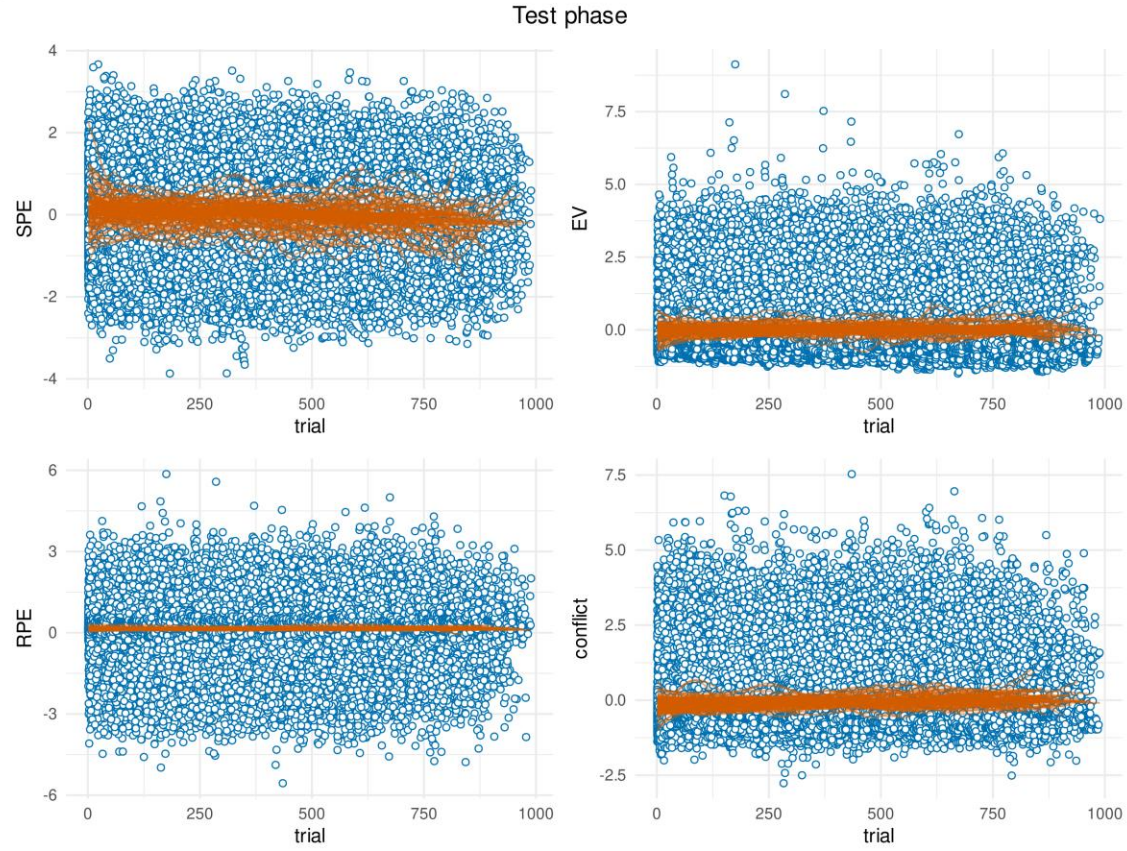


**Fig S10 Temporal evolution of successor-derived regressors during the test phase.** Blue dots represent regressor values of individual trials as assigned under the maximum a-posteriori fit for each participant. Trials are shown in the order they were experienced by the participants. Orange lines are generalized additive models fitted to the evolution of the regressor values over trials, separately for each participant.

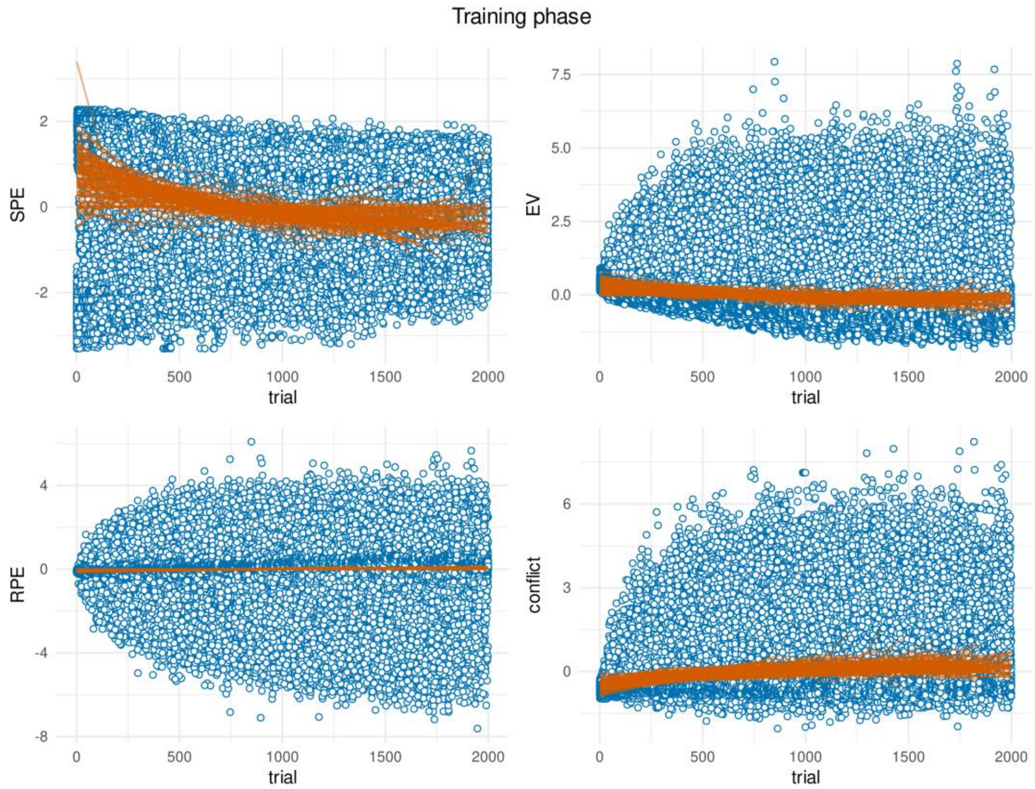


**Fig S11 Temporal evolution of successor-derived regressors during the training phase.** Blue dots represent regressor values of individual trials as assigned under the maximum a-posteriori fit for each participant. Trials are shown in the order they were experienced by the participants. Orange lines are generalized additive models fitted to the evolution of the regressor values over trials, separately for each participant.

### MODULARITY MEASUREMENT

Our "balanced" action-outcome mapping yields "preferred" and "non-preferred" transitions, and also a subset of transitions associated with a "wide" node (see main text methods and Fig. 9). We speculated that "preferred" transitions may be more frequent and thus associated with lower state prediction error, whereas "wide" transitions may be less frequent and therefore associated with higher state prediction error. However, conditional upon selecting a specific action, each transition is always equally likely. Fig. S12A shows state prediction errors associated with a subset of "wide trans" and "preferred" transitions ending in the same state. It can be seen the successor representation model is successfully conditioned on actions, since these different transition types yield identical state prediction errors. Additionally, Fig. S12B shows state prediction errors associated with the "within" and "between" transitions. Neither of these transitions end in a "wide" node, and end in the alternative outcome to a "preferred" transition (main text Fig. 9). With respect to the action-outcome contingencies, they are thus identical. Their only difference is that one maps between communities, while the other maps within communities. The large discrepancy of state prediction error between these transition types provide strong evidence that our successor representation model successfully captures global surprise signals.

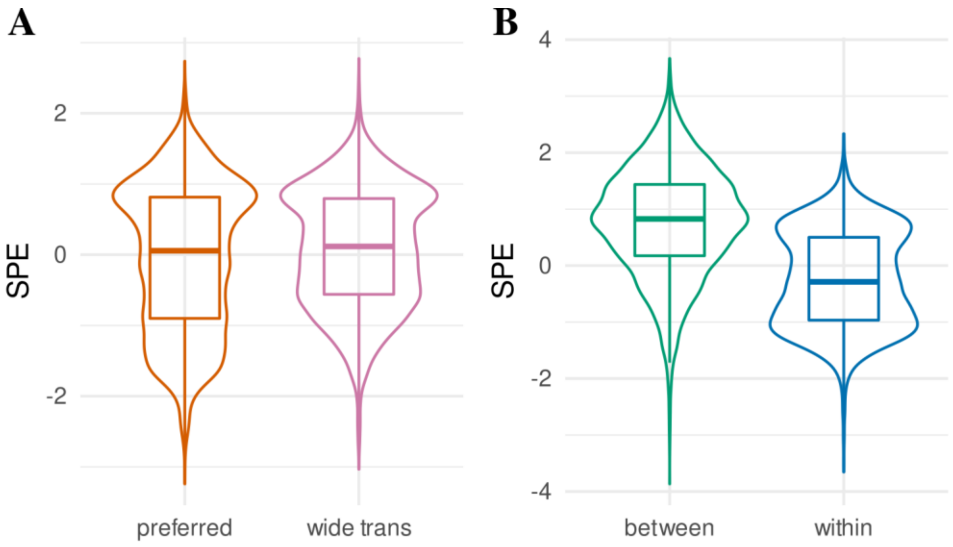


**Fig S12 State prediction errors for different transition types.** Distributions of state prediction errors (SPE) as yielded by the successor representation model of response times (maximum a-posteriori fit for each participant), grouped by transition types associated with the "balanced" action-outcome mapping (colored as in main text Fig. 9). (**A**) We predicted preferred transitions should yield lower SPE, since they are overall more frequent, while transitions out of the wide node should yield increased SPE, since they are overall less frequent. However, when conditioned on the original state, each transition is always equally likely. No obvious difference is present between these two transition types, indicating the successor representation model learned transitions specific for different states and actions, as opposed to marginally over the entire task space. (**B**) Even though transitions between and within communities (colored as in main text Fig. 9) should have identical overall frequency and are never "preferred", the successor representation model of response times assigns higher SPE to the transitions between communities.

#### A SYMMETRIC ACTION-OUTCOME MAPPING

An inelegant aspect of our task design is the removal of certain edges and the duplication of others across both available actions. It is in fact possible to design action-outcome mappings that do not remove or duplicate edges (e.g. Fig. S13A), and yields desired node visitation statistics under both a random walk and a one-directional policy (Fig. S13B). However, when comparing an optimal model-based policy with the explicit hierarchical policy, we can see there is only one room in each wing where the predictions from these models consistently differ (Fig. S13C). In the figure this would correspond to room 2 when taking action i, and room 4 when taking action ii (Fig. S13A). Crucially, the room where this "antirotation" becomes optimal is directly reached by transitioning away from the boundary room connecting to the goal community, when following the rotational action. For example, when in room 1 and taking action i, it is possible to transition to room 4. This is the room where the antirotation is optimal, and the participant should switch to action ii (which has a 50% chance of returning to room 1). However, they might do so more because they identified a failure to make the desired transition into the goal community, instead of having learned that specifically room 4 allows for a more optimal antirotational action. Similarly, if participants learn the heuristic that actions are reversible, they might reverse their action to reach this boundary state again, even though they did not actually learn a detailed model of the museum layout. This could be overcome by excluding trials where room 4 was preceded by room 1 (i.e., exclude trials where the room with an optimal anti-rotation was preceded by a room at the boundary connecting to the goal community).

However, this would come at the cost of statistical power, as it is likely 50% (and even more if participants behave optimally) of trials where the room with an optimal anti-rotation is reached following the boundary room connecting to the goal community. Notably, any action-outcome mapping that would i) maintain all transitions of the original graph as shown in main text Fig. 1A, and ii) not allow for reciprocal transitions given the same action, would display this potential confound. Crucially, participants' sensitivity to lower-level task structure as revealed in main text Fig. 2G are not subject to these limitations given the balanced action-outcome mapping.

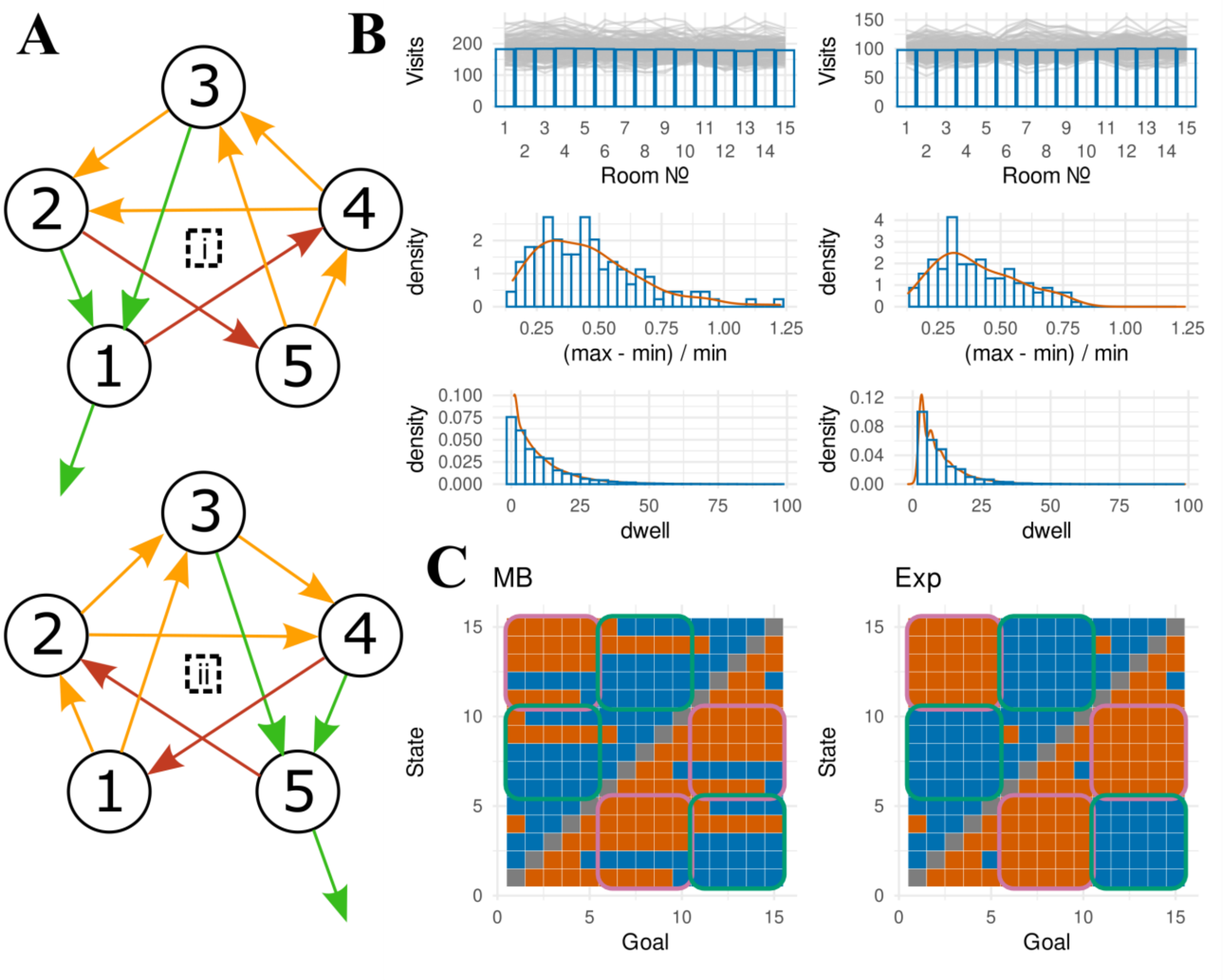


**Fig S13 A symmetric action-outcome mapping.** (**A**) Action-outcome mappings that are symmetric across the two actions (i, anticlockwise rotation; ii, clockwise rotation). Green arrows correspond to outcomes in agreement with the desired rotation, red arrows go against the desired rotation, and orange arrows do not yield progress with respect to the rotation. (**B**) Node visits (top), visit asymmetry (middle), and within-community dwell time (bottom) for the random policy (left) and one-directional policy (right). Further information see description of Fig. S4. (**C**) Chosen actions following a model-based (MB, left) or explicit hierarchical (Exp) policy. Further information see description of Fig. S5.

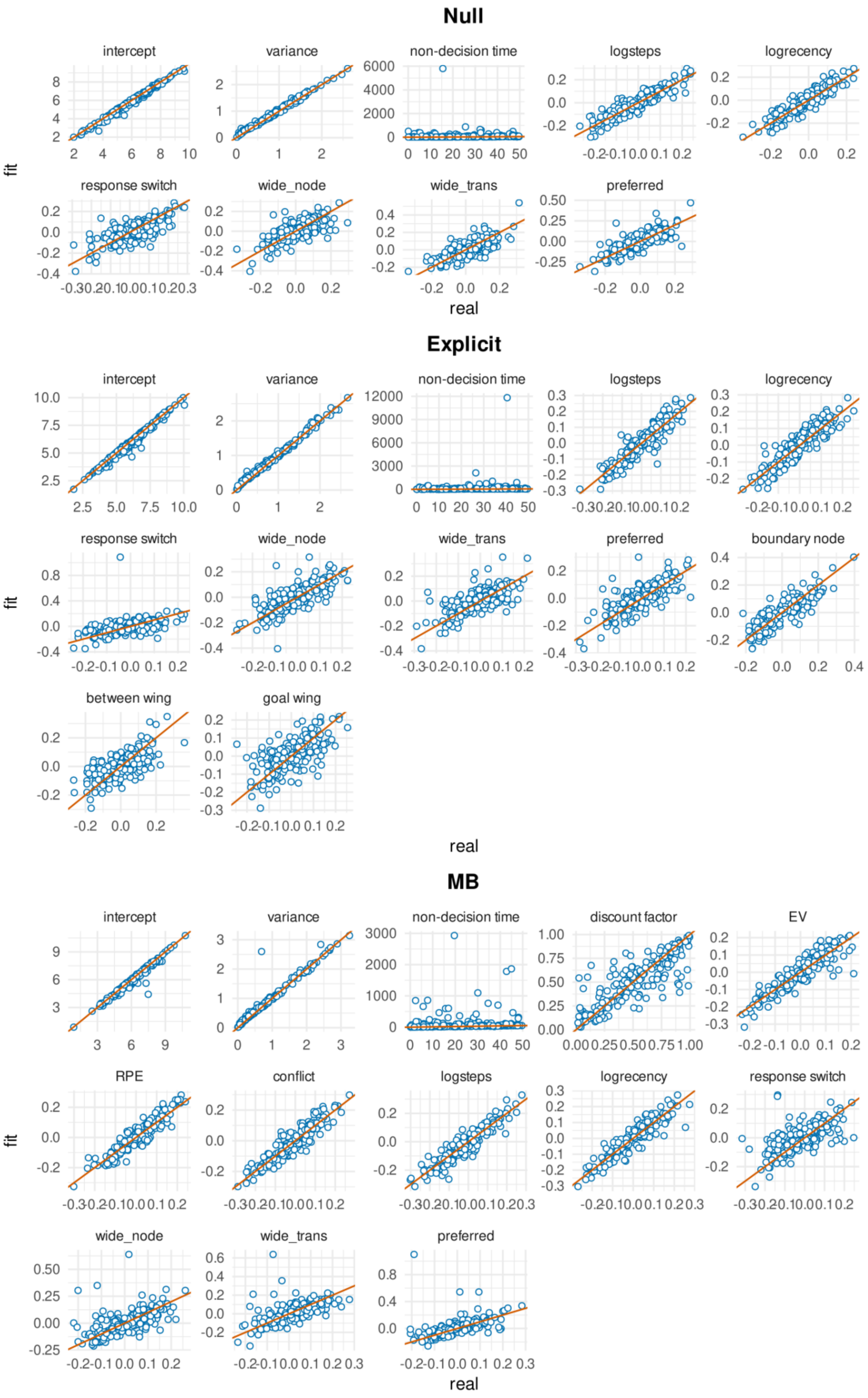

Null
intercept
variance
non-decision time
logsteps
logrecency
response switch
wide_node
wide_trans
preferred
fit
real
Explicit
intercept
variance
non-decision time
logsteps
logrecency
response switch
wide_node
wide_trans
preferred
boundary node
between wing
goal wing
fit
real
MB
intercept
variance
non-decision time
discount factor
EV
RPE
conflict
logsteps
logrecency
response switch
wide_node
wide_trans
preferred
fit
real

**Fig S13 Response time parameter recovery for the null, explicit hierarchical, and model based models.** Ground-truth generated parameters (x-axis) plotted against posterior mean of the fitted parameters (y-axis).
