## Supplementary Materials 2: Control model fits for "The successor representation subserves hierarchical abstraction for goal-directed behavior"

### S2 Appendix: Control model fits

#### CHOICE MODELS

The reported choice models involve the estimation of a discount factor. While the parameter recovery reported in Appendix S1 (Fig. S8) shows good recovery of the discount factor, one could imagine the discount factor is a ‘shallow’ parameter, unreasonably penalizing the successor representation and the model-based models for higher complexity, which might have driven the recovery of the explicit hierarchical model (which does not have a discount parameter). If the parameter were indeed shallow and different estimated values would be inconsequential for the likelihood of the model, approximate leave-one-out cross-validation should take this into account, since it only penalizes models proportional to the ‘diffusiveness’ of predictions based on the posterior distribution (instead of the number of parameters, such as typically used for AIC or BIC). There is thus not much reason to believe the increased model complexity associated with the discount factor unreasonably penalized our more complex models, leading to the significant recovery of the explicit hierarchical model. However, to confirm this inference, we re-fitted the model based and successor representation models of choice behavior for each participant, fixing the discount factor value to the posterior mean as estimated from their corresponding response time model fit. Fig. S15 shows the posterior model probabilities associated with this set of models. This analysis confirms a significant preference for the explicit model (BOR < $10^{-28}$, pxp = 1) that survives binary comparisons (all BOR < $10^{-7}$, all pxp = 1).

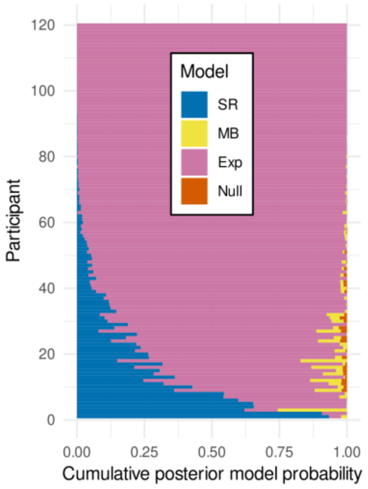


**Fig S15 Posterior model probabilities for the choice models with fixed discount factors.** Horizontally stacked bar charts reflect the posterior model probabilities for the four different choice models. Participants were sorted by posterior model probability of the explicit hierarchical model for clarity.

#### RESPONSE TIME MODELS

While our choice models are arguably simple logistic regressions, our response time models are relatively complex generalized linear models with many independent variables. Here, we aim to justify our modeling choices reported in the main text and show that alternative specifications do not yield significantly different results.

##### Original preregistration

Most crucially, the modeling reported in the main text deviated from our preregistration (https://osf.io/n2jcz/) in three significant ways (see section "control methods" below for preregistered model specification). Firstly, we specified the temporal difference updating target as the outcome state, while we later realized the correct updating target should be the original state (see main text discussion). Secondly, we did not specify nuisance regressors associated with the 'preferred' action-outcome transitions in the preregistration. We believe there are principled reasons to include these regressors, namely because it is reasonable to expect that they might influence the response times. Including these nuisance regressors should only increase the uncertainty about the variables of theoretical interest, thereby making any significant results more robust. Thirdly, we altered the specific implementation of the successor representation to be more similar to the state-action formulation as reported by (1,2), instead of maintaining separate successor matrices based on the actions selected, as we originally proposed. We believe that this change better adheres to common practice as developed in this earlier research. To confirm that these modifications did not induce major changes to the results, here we report results from the exact preregistered analysis protocol. Fig. S16 shows the resulting posterior model probabilities and parameters. These results confirm a significant preference for the successor representation irrespective of the changes made to the model (BOR = 0.005, pxp = 0.997). Nevertheless, the binary comparisons for this model specification are less robust, yielding significance for a binary comparison with the null model (BOR = 0.003, pxp = 0.998) but not with the model based (BOR = 0.064, pxp = 0.968) and explicit hierarchical (BOR = 0.244, pxp = 0.878) models.

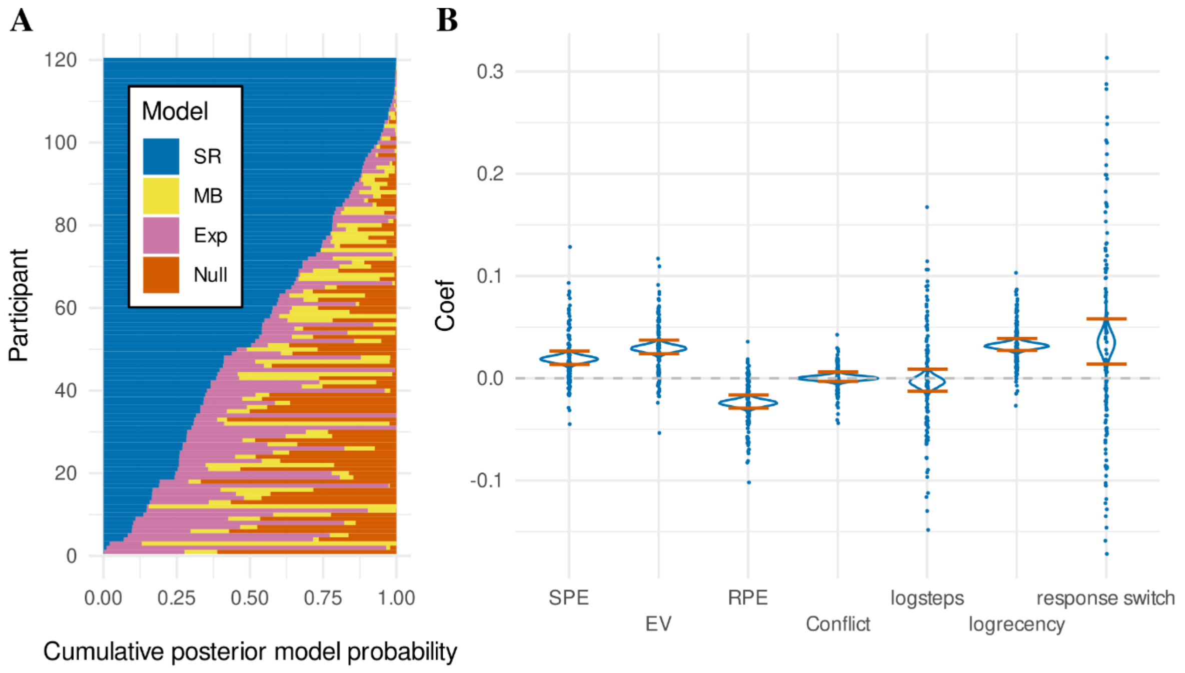

**Fig S16 Preregistered response time analysis.** (**A**) Posterior model probabilities for every participant presented as horizontal stacked bar charts, sorted by successor representation (SR) for interpretability. (**B**) Regression coefficients of the successor representation model. Density plots show the full posterior distributions of the population means, with orange lines indicating the 95% HDI. Dots represent posterior means of individual participant (random) effects. SPE refers to the successor prediction error, EV to expected value, and RPE to reward prediction error.

The preregistered fit of the successor representation confirms a significant slowing effect for larger state prediction errors ("SPE") ($M$ = 0.018, $HDI_{95\%}$ = [0.012, 0.025], $ER_+ = \infty$). It also confirms significant slowing for higher expected values ("EV") ($M$ = 0.029, $HDI_{95\%}$ = [0.022, 0.036], $ER_+ = \infty$) and significant speeding for larger reward prediction errors ("RPE") ($M$ = -0.024, $HDI_{95\%}$ = [-0.031, -0.018], $ER_- = \infty$). Similar to our main reported analysis, the coefficient for the conflict regressor appears quite centered around 0 and does not show evidence for any effect ("Conflict") ($M$ = 0.000, $HDI_{95\%}$ = [-0.004, 0.005], $ER_+$ = 1.017).

With respect to the nuisance regressors, similar to the main reported analysis, the preregistered fit does not find any evidence that participants speed up or slow down based on the (log-transformed) number of trials on the current miniblock ("logsteps") ($M$ = -0.003, $HDI_{95\%}$ = [-0.014, 0.008], $ER_-$ = 2.779). The preregistered fit confirms a significant effect of response slowing for rooms based on (log-transformed) recency ("logrecency") ($M$ = 0.032, $HDI_{95\%}$ = [0.026, 0.038], $ER_+ = \infty$). The preregistered fit also confirms a significant slowing for trials where the participants switched their response key ("response switch") ($M$ = 0.035, $HDI_{95\%}$ = [0.012, 0.057], $ER_+$ = 859.215).

#### State-based successor representation

The successor representation model we presented in the main text considers not only task states, but also the actions selected by the participant, which is required for selecting actions in tasks with probabilistic outcomes. The implementation permitted modelling choices based on the successor representation and defining choice conflict for each trial. However, the successor representation model did not explain the empirical choice data as well as the competing models did, nor did the conflict regressor significantly explain response times. Speaking to this question, neuroscientific literature has proposed that place cell maps observed in the hippocampus may reflect successor-like predictive relationships (3). Crucially, this formulation, as well as the original formulation of the successor representation by (4), only takes into account relationships between states (i.e., marginalized over actions). This formulation could, in principle, explain all of our significant results. Fig. S17A shows the results of the response time model comparison when we include a purely state-based successor representation (see "control methods" for model specification). We also include the original state-action successor representation, and confirm this one is overall preferred (BOR < $10^{-8}$, pxp = 0.995). When inspecting binary comparisons between the state-based successor representation and all other models, a significant preference emerges compared to the null model (BOR = 0.014, pxp = 0.993) and the model-based model (BOR = 0.010, pxp = 0.995). However, a binary comparison with the explicit model does not reach significance (BOR = 0.099, pxp = 0.951). Interestingly, a binary comparison with the original state-action successor representation shows the models perform very comparably, with a slight numerical preference for the original state-action formulation (BOR = 0.831, pxp = 0.577), confirming that both models capture a lot of the same variance.

Inspecting the slopes of the regressors of the response time state-based successor representation (Fig. S17B) confirms a slowing effect for larger state prediction errors ("SPE") ($M$ = 0.035, $HDI_{95\%}$ = [0.026, 0.044], $ER_+ = \infty$). It also confirms significant slowing for higher expected values ("EV") ($M$ = 0.032, $HDI_{95\%}$ = [0.025, 0.039], $ER_+ = \infty$) and significant speeding for larger reward prediction errors ("RPE") ($M$ = -0.020, $HDI_{95\%}$ = [-0.027, -0.014], $ER_- = \infty$). With respect to the nuisance regressors, we again find inconclusive evidence for any influence of the (log-transformed) number of steps of the current miniblock ("logsteps") ($M$ = -0.003, $HDI_{95\%}$ = [-0.014, 0.008], $ER_-$ = 2.693). Rooms that were encountered longer ago for the last time again yielded significant slowing ("logrecency") ($M$ = 0.028, $HDI_{95\%}$ = [0.023, 0.033], $ER_+ = \infty$). Again, trials where the participant switched the keyboard response to the other key were executed significantly slower ("response switch") ($M$ = 0.030, $HDI_{95\%}$ = [0.007, 0.051], $ER_+$ = 275.817).

Similar to the results reported in the main text, nuisance regressors controlling for the "preferred" transitions yielded inconclusive evidence. According to the state based model, participants are significantly slower following more expected transitions, which occur when both actions can transition to the same room ("preferred") ($M$ = 0.019, $HDI_{95\%}$ = [0.011, 0.027], $ER_+ = \infty$). This effect is in the same direction as that of the main state-action model, which did not reach the threshold for a robust effect, whereas it does so for the state-based model. However, contrary to the state-action model reported in the main text, the state-based model yields a trend for participants to be slower in rooms that have more possible outcomes ("wide node", which have no "preferred" transitions) ($M$ = 0.003, $HDI_{95\%}$ = [-0.005, 0.011], $ER_+$ = 3.258) and in the transitions following these rooms ("wide trans") ($M$ = 0.007, $HDI_{95\%}$ = [-0.001, 0.015], $ER_+$ = 22.888). Interestingly, the trending effects for the "wide node" and "wide trans" transitions are in the direction that we hypothesized, whereas they went in the opposite direction for the state-action model (see main text section "Response times").

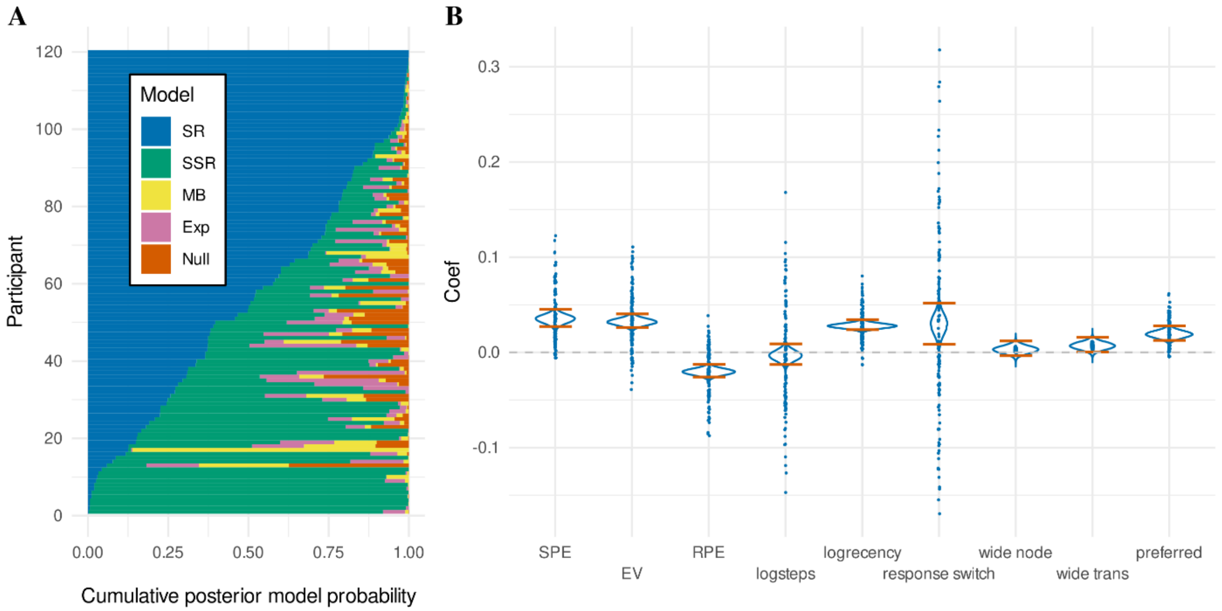


**Fig S17 Response time data including state-based successor representation.** (**A**) Horizontal stacked bar charts for every participant, visualizing the posterior model probability derived from random effects Bayesian model comparison between five cognitive models, "Null" (orange), explicit hierarchical "Exp" (pink), model-based "MB" (yellow), state-based successor representation "SSR" (green), and successor representation "SR" (blue). Participants were sorted by posterior model probability for the "SR" model for interpretability. (**B**) Regression coefficients of the state-based successor representation model. Density plots show the full posterior distributions of the population means, with orange lines indicating the 95% HDI. Dots represent posterior means of individual participant (random) effects. SPE refers to the successor prediction error, EV to expected value, and RPE to reward prediction error.

#### Excluding transitions between wings

Previous research has shown that participants tend to slow down their response times when transitions between stimuli across two different communities are experienced (5–7). The successor representation with a free discount factor parameter is especially suited to capture this pattern of data. It is interesting to see what would happen if we fit the different computational models to the same data, but exclude all trials following a between-community transition from the likelihood. If the successor representation remains the winning model, then this would indicate that it captures more variance than just that which can be described by the community structure of the environment, setting it further apart further from the explicit hierarchical model. In fact, when excluding all between-community transitions from the likelihood during model fitting, the successor representation is still the preferred model of response time data (BOR < $10^{-11}$, pxp = 1) (Fig S18A). This holds for all binary model comparisons between the successor representation and every other model (all BOR < $10^{-4}$, see Methods for multiple comparison correction. All pxp = 1).

Significance tests of all regressors lead to the same conclusion as in the main text, with inconclusive evidence for the influence of (log-transformed) number of trials on the current miniblock ("logsteps") ($M$ = -0.004, $HDI_{95\%}$ = [-0.015, 0.007], $ER_-$ = 2.886), significant slowing for less (log-transformed) recent rooms ("logrecency") ($M$ = 0.029, $HDI_{95\%}$ = [0.024, 0.035], $ER_+ = \infty$), and significant slowing when participant changed their response key ("handswitch") ($M$ = 0.034, $HDI_{95\%}$ = [0.013, 0.055], $ER_+$ = 1094.890). Similarly, we find inconclusive evidence that participant slow down for the node with 2 unique transitions per response ("wide node") ($M$ = 0.004, $HDI_{95\%}$ = [-0.004, 0.012], $ER_+$ = 4.710) or in transitions that follow this node ("wide trans") ($M$ = 0.005, $HDI_{95\%}$ = [-0.003, 0.013], $ER_+$ = 6.416). We find significant slowing for responses following a "preferred" transition ($M$ = 0.020, $HDI_{95\%}$ = [0.011, 0.028], $ER_+ = \infty$).

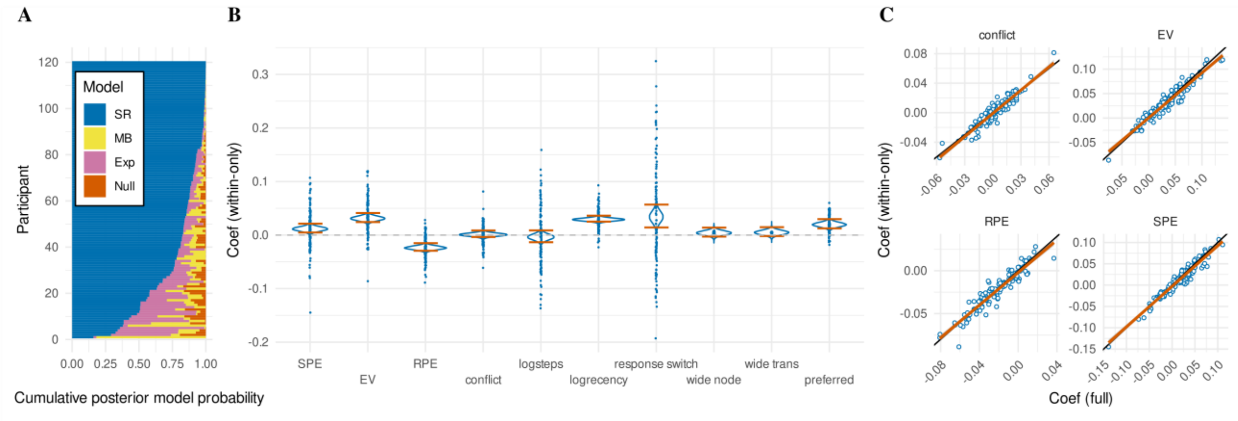


**Fig S18 Models fits for within-wing transitions only.** (**A**) Horizontal stacked bar charts for every participant, illustrating the posterior model probability derived from random effects Bayesian model comparison between four cognitive models, "Null" (orange), explicit hierarchical "Exp" (pink), model-based "MB" (yellow), and successor representation "SR" (blue). Participant data were sorted by posterior model probability for the "SR" model for interpretability. (**B**) Regression coefficients of the successor representation model. Density plots show the full posterior distributions of the population means, with orange lines indicating the 95% HDI. Dots represent posterior means of individual participant (random) effects. SPE refers to the state prediction error, EV to expected value, and RPE to reward prediction error. See text for definition of remaining terms. (**C**) Correlations between posterior means of regressors of theoretical interest for individual participants, as fitted by the full model (x-axis; see also main text Fig. 3) and the within-wing only model (y-axis). Black lines indicate identical values, and orange lines indicate the best fitted regression lines.

With respect to the regressors of theoretical interest (Fig S18B), we find significant slowing for transitions yielding higher state prediction error ("SPE"), indicating slowing when transitioning between two rooms with more different successor representations (e.g. "deep" rooms vs "boundary" rooms) ($M$ = 0.011, $HDI_{95\%}$ = [0.003, 0.020], $ER_+$ = 304.344). We also find significant slowing for higher expected values ("EV"), indicating that participants respond slower for rooms closer to the current goal ($M$ = 0.031, $HDI_{95\%}$ = [0.023, 0.036], $ER_{+}$ = ∞). We also find significant slowing for transitions with more negative reward prediction errors ("RPE"), consistent with post-error slowing when participants move away from the goal, or conversely, speeding when participants move toward the goal ($M$ = -0.024, $HDI_{95\%}$ = [-0.031, -0.017], $ER_{-}$ = ∞). Finally, we find no conclusive evidence that the regressor for conflict, conceptualized as the difference in expected value between the two different actions, influences response times ("conflict") ($M$ = 0.001, $HDI_{95\%}$ = [-0.005, 0.007], $ER_{+}$ = 1.472). The fact that all comparisons yield the same result is not surprising, given the high correlations between parameter estimates of the within-community model fit, compared to the parameter estimates of the full model fit (Fig S18C). Notably, the estimation of the discount factors is consistent whether between-community transitions are included or not (Fig S19). Since the modularity of the successor representation is a direct function of the value of the discount factor, this indicates that measures of modularity of the successor representation can be estimated robustly even without a quantification of response time slowing following transitions between communities. All the relevant model comparisons and parameter estimations are thus found to be significant in the same direction, regardless of including or excluding transitions between communities. We conclude that information about response time slowing between communities does not provide any information that cannot also be extracted from response time variance within communities.

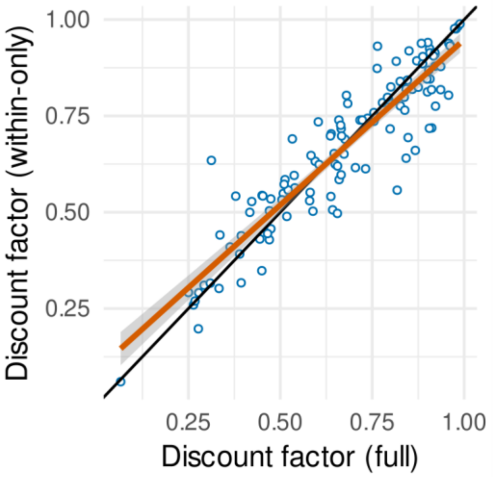


**Fig S19 Discount factor posterior means.** Posterior means of the estimated discount factor for the successor representation models of response times for each participant. The x-axis shows the posterior mean of the discount factor when including transitions between communities, and the y-axis shows the posterior mean of the discount factor when excluding transitions between communities. Black lines indicate identical values, and orange lines indicate the best fitted regression lines.

#### A more equivalent explicit hierarchical model

The explicit hierarchical model considered in the main text, consisted of three non-nuisance regressors, while the successor representation consisted of four. Where the successor representation modeled the expected value of the current room, the explicit hierarchical model encoded a binary value indicating whether the current room was part of the goal wing or not. Where the successor representation encoded a continuous state prediction error, the explicit hierarchical model encoded a binary value indicating whether a transition between two different communities was just made. These two variables provide a proper correspondence between the two models. However, where the successor representation encoded the conflict (implemented as absolute difference between expected value of the two choices), the explicit hierarchical model encoded a binary value indicating whether the current room allowed for a transition between two different communities. To make this correspond better to the actual conflict variable, we here consider a different binary value, that indicates whether the current room lies at the boundary allowing for a transition into or out of the goal wing. These rooms are assigned the highest conflict in the successor representation model. Most crucially, the successor representation encoded the reward prediction error, while the explicit hierarchical model did not have a regressor corresponding to this value. We included an additional variable that was coded as 1 for transitions into the goal community, as -1 for transitions out of the goal community, and as 0 otherwise. Even with this more equivalent definition of the explicit hierarchical model, the successor representation is the preferred model of response time data (BOR $<10^{-10}$, pxp = 1) (Fig S20). This holds when considering a binary comparison between the explicit hierarchical and successor representation models (BOR = 0.001, pxp = 0.999).

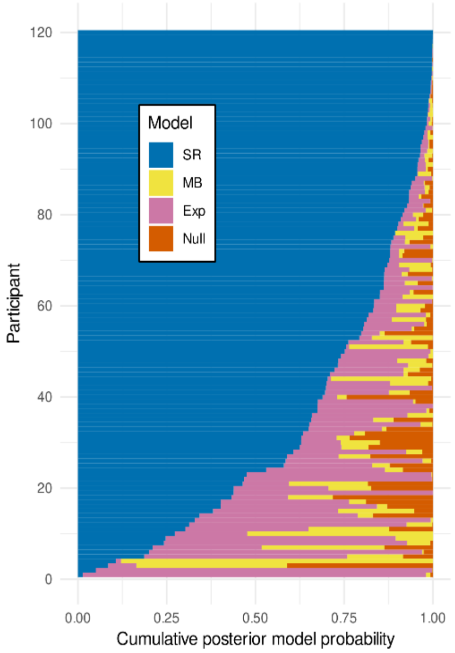


**Fig S20 Model comparison with a more sophisticated explicit hierarchical model.** Horizontally stacked bar charts reflect the posterior model probabilities for the four different response time models. Participants were sorted by posterior model probability of the successor representation model for clarity.

#### JOINT MODELING OF RESPONSE TIMES AND CHOICES

The museum task is entirely self-paced, so choices should be independent from response times: the participants are not pressured to respond quickly before fully evaluating their options. Nevertheless, participants might want to execute their responses quickly in order to finish the experiment as fast as possible, which would be optimal in terms of the average reward per unit of time as opposed to the total reward obtained in the task. Typically, evidence accumulation models that jointly simulate choices and response times are used to account for such speed-accuracy tradeoffs. Joint modeling also allows choice data and response time data to inform each other during parameter estimation, which is especially relevant for the model-based and successor representation models, which directly share the discount factor parameter between the choices and the response times. Regardless, we specifically hypothesized that choices and response times should be best explained by different models, hence our main approach modeled them separately. In this section we report results from joint models instead, to confirm that choices and response times are indeed better explained by different models, even under the assumption (during parameter estimation) that they are derived from the same model. Note that we elected not to use evidence accumulation models for this purpose, as such models introduce significant complexity during estimation and we are not specifically interested in questions regarding the speed-accuracy tradeoff.

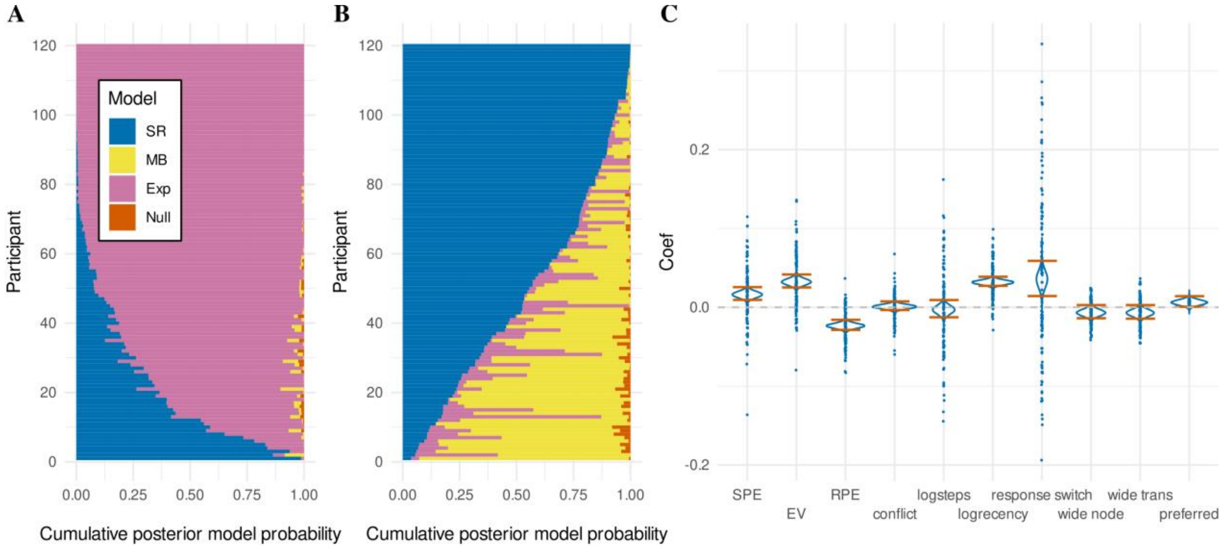


**Fig S21 Joint modeling of choice and response time data.** (**A**) Horizontal stacked bar charts for every participant, visualizing the posterior model probability of the choice data, derived from random effects Bayesian model comparison between four cognitive models, "Null" (orange), explicit hierarchical "Exp" (pink), model-based "MB" (yellow), and successor representation "SR" (blue). Participants were sorted by posterior model probability for the "Exp" model for interpretability. (**B**) As in A, but for the response time data. (**C**) Regression coefficients of the joint successor representation model, with respect to response time data. Density plots show the full posterior distributions of the population means, with orange lines indicating the 95% HDI. Dots represent posterior means of individual participant (random) effects. SPE refers to the successor prediction error, EV to expected value, and RPE to reward prediction error.

Joint modeling confirms the explicit hierarchical model is the preferred model for explaining choice data (BOR < $10^{-26}$, pxp = 1) (Fig. S21A). All binary comparisons remain significant (all BOR < $10^{-4}$, all pxp = 1). The estimated regression coefficient for the explicit model is identical to the one reported in the main text (Fig. 2B), because the model specification is identical, thus we do not report this again.

Joint modeling confirms the successor representation is the preferred model for explaining response times (BOR < $10^{-8}$, pxp > 0.999) (Fig. S21B). Binary comparisons remain significant with a preference for the successor representation model compared the null model (BOR < $10^{-8}$, pxp = 1) and the explicit model (BOR = 0.003, pxp = 0.997). However, the preference for the successor representation over the model-based model is no longer significant (BOR = 0.278, pxp = 0.861). This can be explained by the reduced quality of the discount factor estimation, which is now 'contaminated' by information from the choice data, which in fact appears not to be generated according to a successor representation (or a model-based model). The estimated regression coefficients (Fig. S21C) confirm a significant slowing effect for larger state prediction errors ("SPE") ($M$ = 0.016, $HDI_{95\%}$ = [0.008, 0.024], $ER_{+} > 2.667*10^{4}$). They also confirm significant slowing for higher expected values ("EV") ($M$ = 0.032, $HDI_{95\%}$ = [0.024, 0.040], $ER_{+} = \infty$) and significant speeding for larger reward prediction errors ("RPE") ($M$ = -0.023, $HDI_{95\%}$ = [-0.030, -0.017], $ER_{-} = \infty$). Similar to our main reported analysis, the coefficient for the conflict regressor appears centered around 0 and does not show evidence for any effect ($M$ = 0.001, $HDI_{95\%}$ = [-0.005, 0.006], $ER_{+}$ = 1.599).

With respect to the nuisance regressors, the joint fit replicates the non-significant effect of the (log-transformed) number of steps taken on the current miniblock ("logsteps") ($M$ = -0.003, $HDI_{95\%}$ = [-0.014, 0.008], $ER_-$ = 2.428). The joint fit confirms significant slowing for rooms based on (log-transformed) recency ("logrecency"), indicating rooms that have not been encountered for longer yield slower response times ($M$ = 0.032, $HDI_{95\%}$ = [0.026, 0.037], $ER_+ = \infty$). The joint fit also confirms significant slowing for trials where the participant switched their response key ("response switch") ($M$ = 0.036, $HDI_{95\%}$ = [0.013, 0.058], $ER_+$ = 1289.323).

Further, the joint fit confirms trending effects for an influence of the action-outcome mapping on response times (main text Fig. 3B). A similar trend as the main model finds participants to be faster in rooms that have more possible outcomes ("wide node", which have no "preferred" transitions) ($M$ = -0.007, $HDI_{95\%}$ = [-0.015, 0.002], $ER_-$ = 17.112) and the transitions following these rooms ("wide trans") ($M$ = -0.007, $HDI_{95\%}$ = [-0.016, 0.002], $ER_-$ = 17.972). Meanwhile, the joint model replicates a trend for participants to be slower following more expected transitions, which occur when both actions can transition to the same room (Fig. 1C) ("preferred") ($M$ = 0.006, $HDI_{95\%}$ = [-0.001, 0.013], $ER_+$ = 26.797).

#### Posterior predictive check

As shown in main text Fig. 4, in general our successor representation model fits the true response times well. Not only does the model mostly capture the order of transition types in terms of speed of responding (main text Fig. 4C), it also captures smooth patterns over regressors derived from the successor representation (main text Fig. 4D). However, the model seemed to underestimate response times when participants transitioned into the goal wing (main text Fig. 4C, "into"). Here, we fit and compare two amendments of the model that might account for this finding.

##### Models

Firstly, we considered to replace the signed reward prediction error with an unsigned reward prediction error. The main posterior predictive check revealed that response times seem to increase when reward prediction errors move further away from 0, regardless of whether they are positive or negative (main text Fig. 4D). When including signed reward prediction errors, transitions out of the goal wing will yield a large negative reward prediction error (main text Fig. 4B). Since we estimated a negative regression coefficient for the reward prediction error (main text Fig. 3B), this corresponds to a slowing of response time. This effect likely allows our model to fit the observed pattern in the real data, where transitions out of the goal wings are slower compares to transitions between non-goal wings (main text Fig. 4C). Crucially, these trials both yield large state prediction errors and have very comparable expected values. The reward prediction error is the only significant variable dissociating these trial types. However, transitions into the goal wing yield larger positive reward prediction errors (main text Fig. 4B). The negative regression coefficient then implies responses would speed up when entering the goal wing. This is reflected in our posterior predictions as a speeding up of transitions into the goal community, whereas in the real data these transitions are the ones yielding the slowest response times (main text Fig. 4C). Since the real data seem to imply a slowing effect for absolute reward prediction error, the signed reward prediction error appears as a misspecification of our model yielding the erroneous pattern in this posterior predictive check. Interestingly, even though the model incorporated the sign of the reward prediction errors, inspection of the posterior predictions binned by reward prediction error suggests that the response times in fact increase with absolute reward prediction errors (main text Fig. 4D). This could result from the reward prediction error covarying with the other regressors, for example, because the between-wing transitions elicit large state prediction errors and large positive and negative reward prediction errors, all of which predict slower response times. Hence, even though the model appears to capture the fine-grained pattern of response times over reward prediction error, we fitted a new model that incorporated the absolute reward prediction error (instead of the signed reward prediction error) as a regressor, in order to address the apparent misspecification of the original model with respect to the transitions into the goal wing.

Secondly, we considered adding a regressor that specifically captures moving into the goal wing. Participants could have explicitly used knowledge about the communities to monitor for these transitions. This possibility is akin to setting the relevant bottleneck transition as a "subgoal" in terms of hierarchical reinforcement learning. If true, this possibility should in fact be accounted for by the explicit hierarchical model (which was not the case). However, it is also possible that the participants' response times are influenced primarily from learned successor representations, while also containing an element of hierarchical representation. This latter possibility appears likely, given that the choices are better described by an explicit hierarchical mechanism. We thus fitted an additional successor representation model to the response times that included an additional binary regressor indicating 1 for trials when the participants transitioned into the goal wing and 0 otherwise.

#### Results

A model comparison (Fig. S22A) between the successor representation reported in the main text, the control model with absolute reward prediction errors, and the control model with an additional regressor for subgoals, indicates a non-significant result (BOR = 0.666) but a numerical preference for the model with absolute reward prediction error (pxp = 0.554) over the original (pxp = 0.222) and subgoal (pxp = 0.224) models.

Inspecting the estimated regression coefficients (Fig. S22B) of the absolute reward prediction error model, we confirm a significant slowing effect for larger state prediction error ("SPE") ($M$ = 0.012, $HDI_{95\%}$ = [0.004, 0.020], $ER_{+}$= 417.848) and for higher expected values ("EV") ($M$ = 0.023, $HDI_{95\%}$ = [0.015, 0.031], $ER_{+} = \infty$). Interestingly, we now observe significant slowing effect for absolute reward prediction error ("|RPE|") ($M$ = 0.014, $HDI_{95\%}$ = [0.006, 0.022], $ER_{+} > 2.161*10^{3}$), as opposed to a significant speeding effect for larger positive reward prediction errors as reported by our main analysis. Furthermore, we again find no significant effect of conflict ($M$ = -0.005, $HDI_{95\%}$ = [-0.011, 0.001], $ER_{-}$ = 20.639).

Inspecting the posterior predictions for the absolute reward prediction error model (Fig. S22C), it can be seen that this model properly captures the increased slowing for transitions into the goal community. The response times for all transition types as listed in main text Fig. 4A are now accurately predicted. Interestingly, when inspecting the posterior predictions against the magnitude of the regressors derived from this model, we now see a clear association between slower response times and larger absolute reward prediction errors (Fig. S22D).

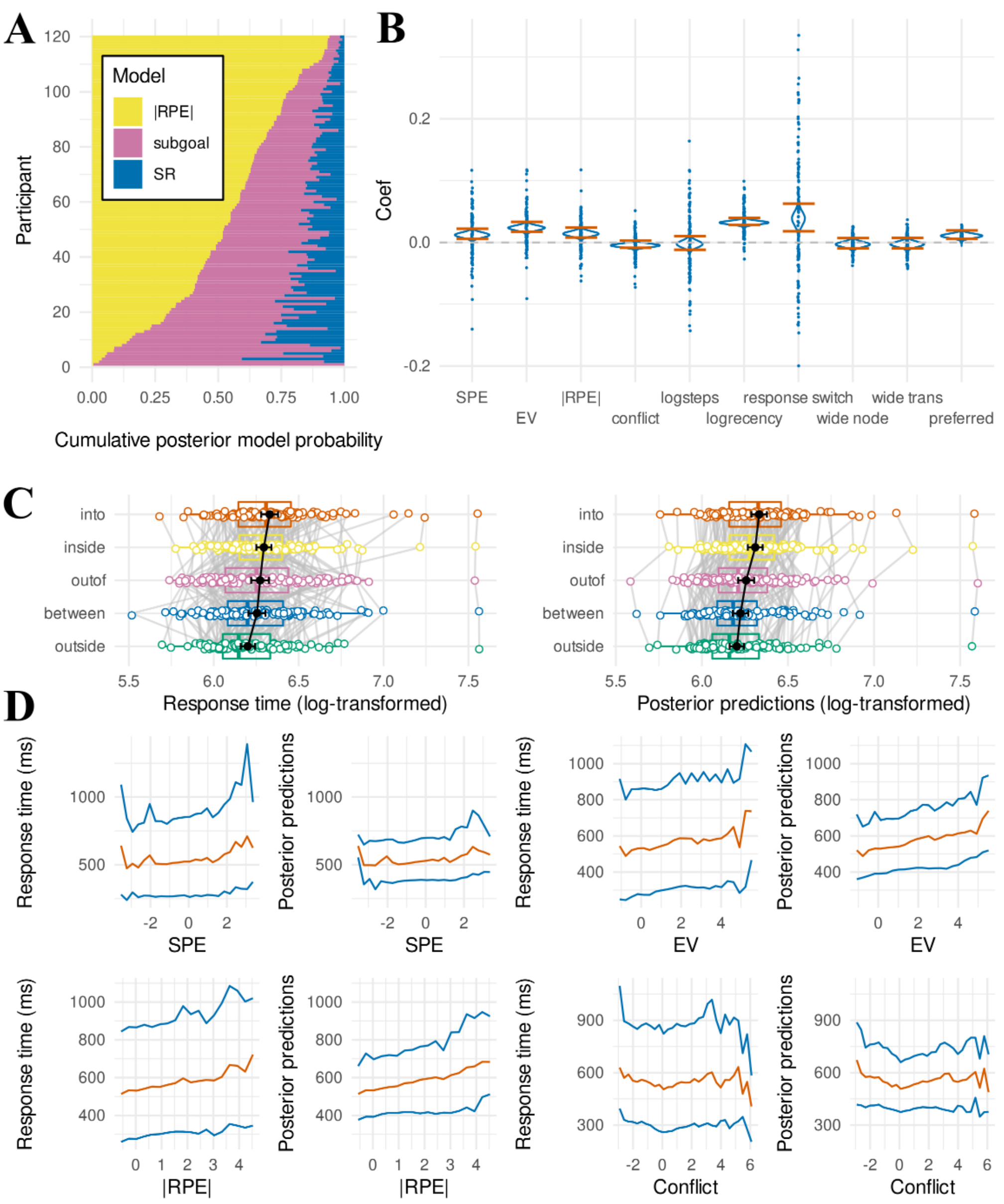


Fig S22 Posterior predictive check of the response time successor representation model with absolute reward prediction errors. (A) Horizontal stacked bar charts for every participant, visualizing the posterior model probability derived from random effects Bayesian model comparison between three cognitive models, the absolute reward prediction error successor representation "|RPE|" (yellow), the successor representation with an additional subgoal regressor "subgoal" (pink), and the main successor representation "SR" (blue). Participants were sorted by posterior model probability for the "|RPE|" model for interpretability (B) Regression coefficients of the successor representation model with absolute reward prediction error. Density plots show the full posterior distributions of the population means, with orange lines indicating the 95% HDI. Dots represent posterior means of individual participant (random) effects. SPE refers to the successor prediction error, EV to expected value, and |RPE| to absolute reward prediction error. (C) Mean (log-transformed) response times (left) and mean posterior predictions of the |RPE| model (right) for each participant for the different transition types as defined in main text Fig. 4A. Population means with their 95% confidence intervals are shown in black. (D) Mean (orange) and 10th and 90th percentile (blue) of the response time distribution for trials binned by regressor values in intervals of 0.3. Binned separately for the different regressors (SPE, EV, |RPE|, conflict). Conditional response time distributions are repeatedly plotted side by side for real data (left) and the full posterior predictive distribution of the |RPE| model (right).

### CONTROL METHODS

#### Preregistered successor representation

The preregistered successor representation indexed separate state-based successor representations for different actions. This representation could be written as $\mathbf{M} \in \mathbb{R}^{|A|\times|S|\times|S|}$. Each entry $\mathbf{M}_{a,s,s'} = \mathbb{E}[\sum_{t=0}^{\infty} \gamma^t \mathbb{I}_{s=s'} | s_0 = s, a_0 = a]$. This can be learned following a SARSA update rule. Starting $\mathbf{M}$ out as $\mathbf{M}[a,:,:] = I_{|S|}$ , an identity matrix for each action $a$, updates follow:

$$\mathbf{M}[a_t, s_t, :] \leftarrow \mathbf{M}[a_t, s_t, :] + \lambda(\mathbb{1}_{s_t} + \gamma \mathbf{M}[a_{t+1}, s_{t+1}, :] - \mathbf{M}[a_t, s_t, :])$$

where $\mathbb{1}_i$ indicates a one-hot vector with a 1 at index *i*.

All other features with respect to deriving the regressors and the specification of priors are the same as for the state-action successor representation, described in the main text methods.

#### State-based successor representation

The state-based successor representation holds no regard for different actions selected by the agent. This representation could be written as $\mathbf{M} \in \mathbb{R}^{|S|\times|S|}$. Each entry $\mathbf{M}_{s,s'} = \mathbb{E}[\sum_{t=0}^{\infty} \gamma^t \mathbb{I}_{s=s'} | s_0 = s]$. This can be learned following a SARSA update rule. Starting $\mathbf{M}$ out as an identity matrix of size $|S|$, updates follow:

$$\mathbf{M}[s_t, :] \leftarrow \mathbf{M}[s_t, :] + \lambda(\mathbb{1}_{s_t} + \gamma \mathbf{M}[s_{t+1}, :] - \mathbf{M}[s_t, :])$$

where $\mathbb{1}_i$ indicates a one-hot vector with a 1 at index *i*.

All other features with respect to deriving the regressors and the specification of priors are the same as for the state-action successor representation, described in the main text methods.

#### Joint model of choices and response times

For the model-based and the successor representation models, the discount factor parameter is shared between the choices and the response time models, which were fitted separately for the models reported in the main text. Joint models combined the choice and response time models in a single Stan program, which considers both choice and response time data. This amounts to simply running a single HMC procedure for the parameters as specified in the main text methods, including both the parameters described under the section “response time model specification” and under the section “choice model specification”, estimating only a single discount factor $\gamma_{im}$. If response times and choices were both generated from the same model with a shared discount factor, this would improve the estimation of the discount factor, and therefore improve the estimation of other parameters, such as the regression coefficients of interest. However, if one of two data streams is not generated from this process, it can introduce bias in the estimation of the discount factor, and therefore in the estimation of the regression coefficients of interest.
